# Parabrachial Opioidergic Projections to Preoptic Hypothalamus Mediate Behavioral and Physiological Thermal Defenses

**DOI:** 10.1101/2020.06.23.167619

**Authors:** Aaron J. Norris, Jordan R. Shaker, Aaron L. Cone, Imeh B. Ndiokho, Michael R Bruchas

**Affiliations:** Department of Anesthesiology, Washington University School of Medicine, St. Louis MO, 63110, USA; Medical Scientist Training Program, University of Washington, Seattle, WA 98195, USA; Center for the Neurobiology of Addiction, Pain and Emotion, Departments of Anesthesiology and Pharmacology, University of Washington, Seattle, WA 98195, USA

**Author notes:** Authors contributed equally (Aaron J. Norris, Jordan R. Shaker).

## Abstract

Maintaining stable body temperature through environmental thermal stressors requires detection of temperature changes, relay of information, and coordination of physiological and behavioral responses. Studies have implicated areas in the preoptic hypothalamic area (POA) and the parabrachial nucleus (PBN) as nodes in the thermosensory neural circuitry and indicate the opioid system within the POA is vital in regulating body temperate. In the present study we identify neurons projecting to the POA from PBN expressing the opioid peptides Dynorphin (Dyn) and Enkephalin (Enk). We determine that warm-activated PBN neuronal populations overlap with both Dyn and Enk expressing PBN populations. We demonstrate that Dyn and Enk expressing neurons are partially overlapping subsets of a glutamatergic population in the PBN. Using optogenetic approaches we selectively activate projections in the POA from PBN Dyn, Enk, and VGLUT2 expressing neurons. Our findings demonstrate that Dyn, Enk, and VGLUT2 expressing PBN neurons are critical for physiological and behavioral heat defense.

## Introduction

Maintaining body temperature in the face of changing environmental conditions is a core attribute of mammals, including humans, and is critical for life. Achieving a stable body temperature requires information about the temperature of the periphery and environment to be integrated to drive physiological and behavioral programs to defend the core temperature (Jessen, 1985). Physiological parameters modulated to maintain temperature include thermogenesis (utilization of brown adipose tissue (BAT), shivering), changes in circulation (vasodilation and vasoconstriction), and evaporation (Cabanac, 1975). Behavioral modifications include selection, when possible, of ambient temperature, altering posture to alter heat loss, and modulation of physical activity level. Responding to thermal challenges involves perception of temperature, encoding the valence of the temperature (ex. too hot), and evoking appropriate physiological responses (Tan and Knight, 2018). Perceptive, affective, and autoregulatory elements may be encoded by overlapping or discrete neuronal circuits. The preoptic area of the hypothalamus (POA) and the parabrachial nucleus (PBN) have been identified as key nodes within the neurocircuitry regulating body temperature. In the report presented here, we identify and delineate the unique roles of genetically defined neuronal populations in PBN projecting to the POA in responding to environmental warmth.

The POA contains neurons critical for integration of information about body temperature and for coordination of responses to thermal challenges to maintain core temperature (Abbott and Saper, 2017, 2018; Tan et al., 2016). Neurons in POA, identified by different genetic markers, can regulate brown adipose tissue (BAT) activation, drive vasodilation, and shift ambient temperature preferences (Tan et al., 2016; Yu et al., 2016). Prior evidence has suggested critical roles for inputs from the PBN to the POA in regulating temperature (Geerling et al., 2016; Miyaoka et al., 1998; Morrison, 2016). The PBN is, however, a highly heterogenous structure with multiple subpopulations known to relay various sensory information from the periphery (thirst, salt-appetite, taste, pain, itch, temperature, etc) (Chiang et al., 2020; Kim et al., 2020; Palmiter, 2018). The studies here examine roles for parabrachial glutamatergic neurons expressing the opioid peptides dynorphin and enkephalin.

Regulation of body temperature requires integration of many inputs across varying time scales, and multiple neuromodulatory neuropeptides may be involved in controlling body temperature. *In vivo* experiments suggest that opioid neuropeptides, as a neuromodulator, may play a critical role in thermal homeostasis. Systemic administration and local infusion into the POA of opioid receptor agonists and antagonists induces changes in body temperature and can impair thermoregulatory control in humans, rats, and other animals (Chen et al., 2005; Ikeda et al., 1997; Spencer et al., 1990). Several studies have implicated opioid receptor signaling within the POA in modulating body temperature, but potential sources for native ligands remain to be identified. (Baker and Meert, 2002; Clark, 1979). Activation of *mu* or *kappa* receptors in the POA can drive opposing effects on body temperature. A recent study indicated that neurons in PBN expressing *Preprodynophin,* which is processed to Dynorphin (Dyn), the endogenous ligand for the Kappa Opiate Receptor (KOR), are activated by ambient warmth (Chavkin et al., 1982; Geerling et al., 2016). PBN neurons expressing the endogenous *mu* and *delta* opioid receptor ligand, enkephalin (Enk), have not been examined in relation to how they may regulate temperature.

In this study we used a series of modern anterograde and retrograde viral approaches to determine the connection of PBN neurons expressing Dyn (Dyn+) and Enk (Enk+), to the POA (Henry et al., 2017). We delineate the overlap between these peptide expressing neuronal populations with warm-activated PBN neurons. We identify subsets of Dyn and Enk expressing neurons that project to POA from the PBN. We then combine optogenetic and chemogenetic tools with Cre driver mouse lines to determine the causal roles of PBN neurons that project to the POA in mediating physiological and behavioral responses to thermal challenge. Here we also examine potential roles of opioid receptor mediated behaviors in both Dyn+ and Enk+ PBN-POA projections. Here we report that glutamatergic, Dyn+, and Enk+ neuronal populations projecting from PBN to POA initiate physiological and behavioral heat defensive behaviors. Chemogenetic inhibition of glutamatergic PBN neurons blocks vasomotor responses to thermal heat challenge. The studies reported here provide new insights into the thermoregulatory properties of parabrachial neuropeptide-containing projections to the hypothalamus in homeostatic and metabolic behavior.

## Results

### Ambient warmth activates Dyn+ and Enk+ neurons in PBN

Effects of *mu* and *kappa* receptor signaling on body temperature have been described and mRNA for *preprodynorphin* and *preproenkephalin* has been reported to be expressed in the PBN (Baker and Meert, 2002; Chen et al., 2005; Clark, 1979; Engstrom et al., 2001; Hermanson and Blomqvist, 1997; Hermanson et al., 1998). To examine if PBN neurons expressing dynophin or enkephalin opioid neuropeptides are activated by ambient warmth, we exposed mice to ambient warmth (38°C) or room temperature (21-23°C) for four hours prior to preparation of brain for fos staining. We performed immunohistochemistry (IHC) on collected brains sections containing the PBN with antibody directed against c-Fos (anti-c-Fos) to examine induction of c-Fos expression as a marker of neuronal activation (Sheng and Greenberg, 1990). Consistent with recent reports, we observed induction of c-Fos expression in the lateral PBN (LPBN) (**Figure 1B-C**) (Geerling et al., 2016). In brain sections from warm exposed mice (n=8) compared to room temperature controls (n=4), c-Fos staining revealed a robust and significant (p=0.003) increase in mean ±SEM number of neurons positive for c-Fos expression in the LPBN per brain: 265.8 ±41.9 in warm exposed mice compared to 23.2 ±4.0 in room temperature controls (**Figure 1C**). Cells in LPBN, lateral to superior cerebellar peduncle, in sections corresponding −5.0 to −5.4 caudal to bregma were counted. In brains from recombinase reporter mice (Ai14) crossed to Dyn-Cre (Ai14xDyn-Cre) or Enk-Cre (Ai14xEnk-Cre) lines, tdTomato was robustly expressed in LPBN indicating expression of *preprodynorphin* (Al-Hasani et al., 2015; Francois et al., 2017; Krashes et al., 2014; Madisen et al., 2010) and *preproenkephalin* (Francois et al., 2017) in LPBN neurons (**Figure 1D-E**). Cells expressing tdTomato in Dyn-Cre mice (Dyn+) and Enk-Cre mice (Enk+) were most abundant in the caudal LPBN. To determine the overlap between warm-activated neurons with Dyn+ or Enk+ cells in LPBN we exposed mice, Ai14xDyn-Cre and Ai14xEnk-Cre, to a warm (38°C) ambient temperature for 4 hours prior to harvesting brains and used IHC on sections with anti-c-Fos. In the LPBN of Ai14xDyn-Cre mice, we found that a mean ±SEM of 81% ±2.5 of the cells positive for c-Fos staining were also positive for tdTomato expression (n=4 animals, 1017 cells) (**Figure 1F**). In the LPBN of Ai14xEnk-Cre mice, an average ± SEM of 54% ±4.6 (n=4 animals, 1109 cells) of c-Fos-positive cells in warm-exposed mice were also positive for tdTomato (**Figure 1G**). These data indicate that warmth activated neurons in LPBN co-express the neuropeptides dynorphin and enkephalin.

**Figure 1.**
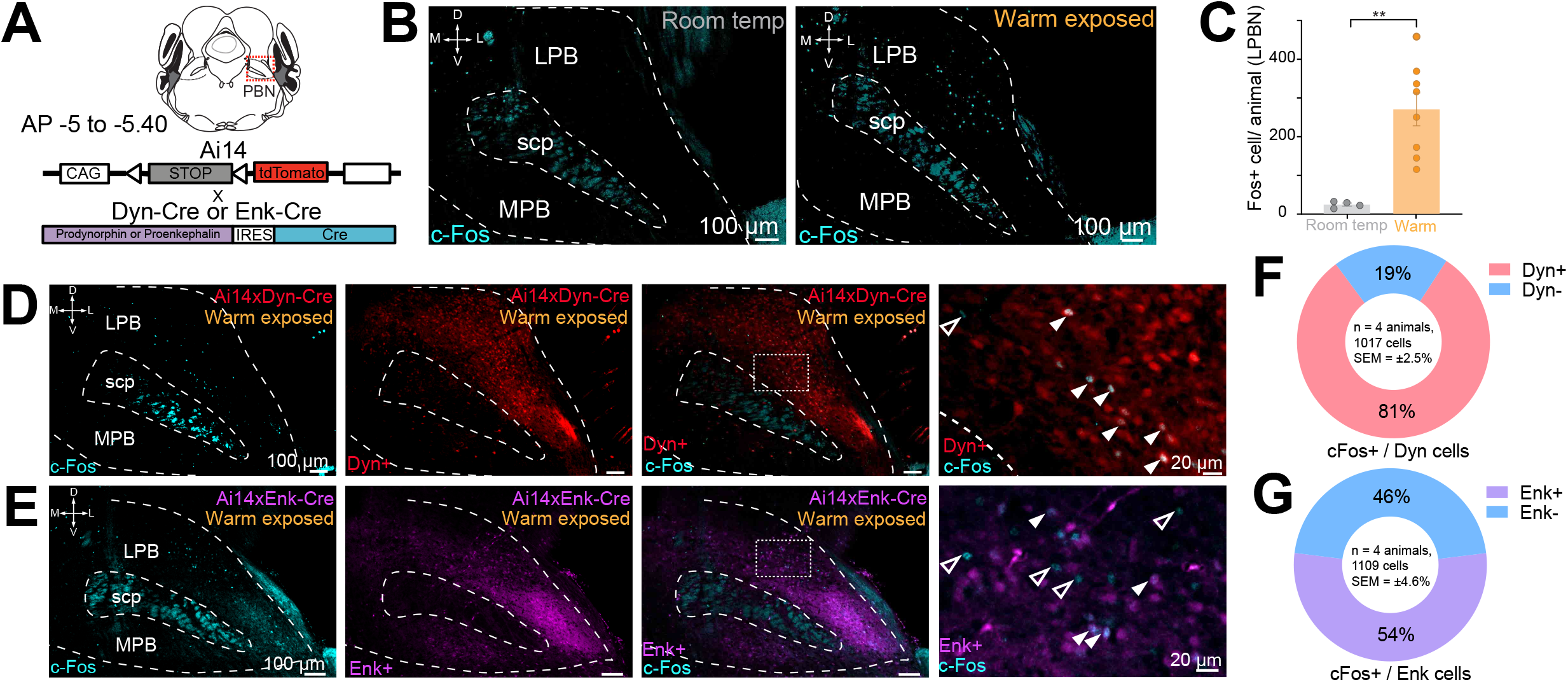
Warm-activated neurons in PBN overlap with Dyn and Enk expression. (**A**) Schematized view of PBN regions analyzed for c-Fos expressing neurons and the genetic cross schemes of Ai14xDyn-Cre/Ai14xEnk-Cre reporter mouse lines used. (**B**) Representative images of brain sections harvested from animals exposed to room temperature or 38°C and probed with anti-c-Fos. Brains from 38°C exposed mice had significantly more neurons in PBN positive for c-Fos staining. (**C**) Quantification of c-Fos positive LPBN neurons per brain. Data are presented as mean ± SEM; n = 4 animals in room temp group, n = 8 animals in warm exposed group; t test, **p < 0.01. (**D**) Representative images of c-Fos labeling (cyan) in Ai14 x Dyn-Cre brains with c-Fos labeling of Dyn+ (red) (filled arrows) neurons and Dyn-(open arrows). (**E**) Representative images of c-Fos labeling in Ai14xEnk-Cre brains with c-Fos labeling of Enk+ (magenta) (filled arrows) and Enk-neurons (open arrows). (**F-G**) Quantification of the overlap of c-Fos staining in Ai14xDyn-Cre and Ai14xEnk-Cre brains demonstrated 81% or 46% of c-Fos cells were also overlapped with tdTomato expression in Ai14xDyn-Cre or Ai14xEnk-Cre brains, respectively.

### Dyn+ and Enk+ PBN neurons project to the ventral medial preoptic area and are VGLUT2+

Next, we used fluorescent *in situ* hybridization (FISH) with target probes for mouse *Preproodynorphin* (pDyn), *Preproenkephalin* (pEnk), and *Slc17a6* (VGLUT2) and examined serial coronal brain sections encompassing the PBN to delineate possible overlapping expression of the neuropeptides. Based on previous studies implicating glutamate in LPBN thermosensory relay neurons (Nakamura and Morrison, 2008, 2010), we hypothesized that the majority of pDyn and pEnk expressing LPBN neurons would be VGLUT2+. Consistent with expression patterns evident in the Ai14xDyn-cre and Ai14Enk-Cre mice, pDyn and pEnk FISH probes labeled neurons in the LPBN (**Figure 2F and G)**. pDyn+ and pEnk+ cells were most abundant in the caudal LPBN. Sections were also co-labeled with VGLUT2 probes with pDyn or pEnk probes. The overlap of cells in PBN labeled with each probe was quantified. A mean ±SEM of 98% ±0.9 (n= 760 cells, n=4 mice) of pDyn labeled cells were positive for VGLUT2 (**Figure 2A, I**). A mean ±SEM of 97% (n= 650 cells, n=4 mice) of pEnk labeled cells were positive for VGLUT2 (**Figure 2B, J**). Surprisingly, a mean ±SEM of 51% ±6.6 (n= 760, n = 4 mice) of PBN neurons positive for pDyn were also positive for pEnk labeling (**Figure 2C, K**). Reciprocally, a mean ±SEM of 58% ±2.3 (n= 650 cells, n=4 mice) of cells labeled by pEnk probes were also labeled by pDyn probes (**Figure 2D, K**). These FISH based experiments indicate that pDyn+ and pEnk+ cells in the PBN express VGLUT2 and are partially overlapping subpopulations of glutamatergic PBN cells.

**Figure 2.**
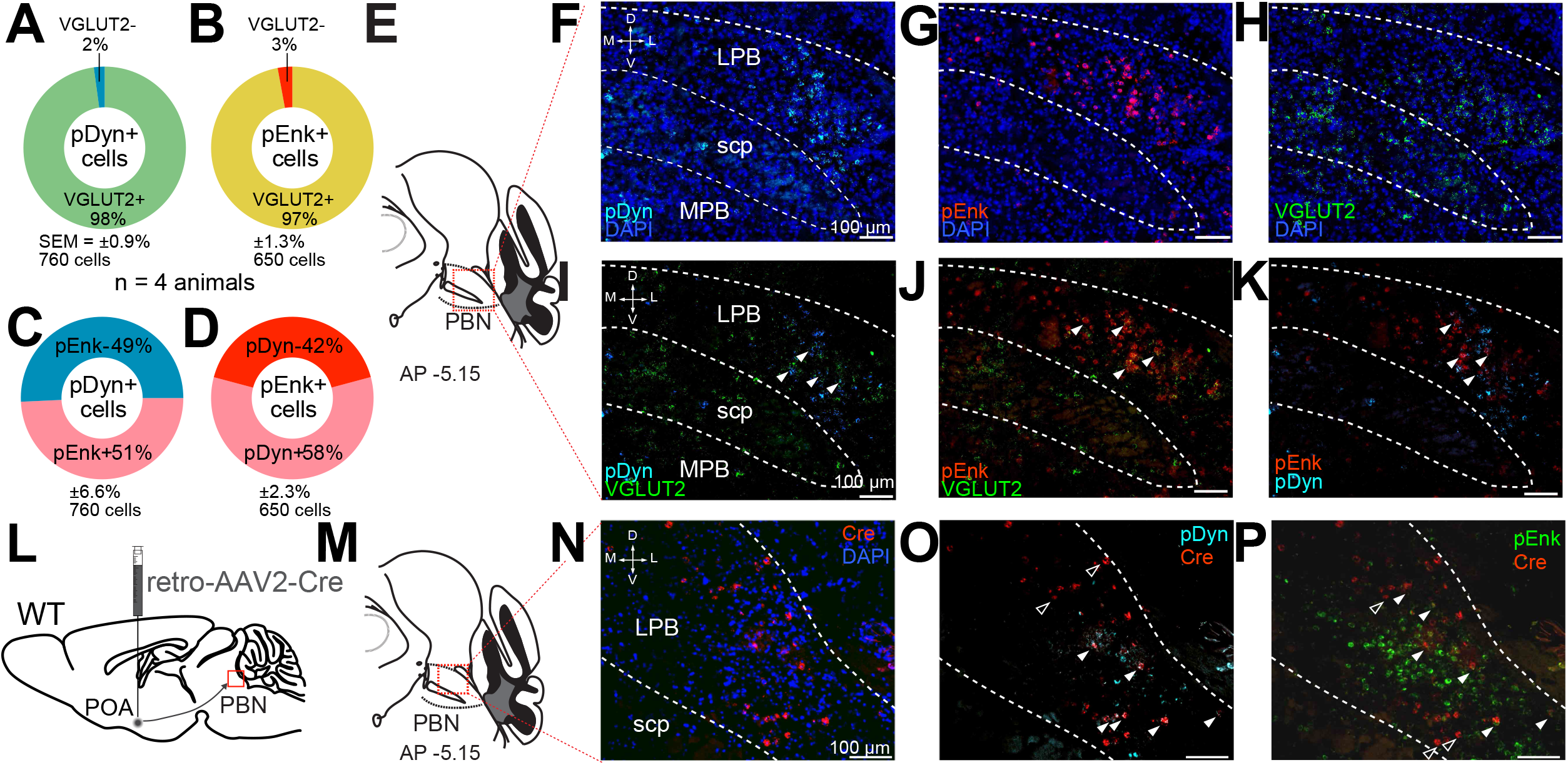
pDyn+ and pEnk+ LPBN neuron populations overlap, are VGLUT2+, and project to the POA. (**A-D**) Quantification of cells labeled with (**A**) pDyn probe (pDyn+) and Slc17a6 (VGLUT2+) probes, or (**B**) pEnk (pEnk+) and Slc17a6 (VGLUT2+) probe, or (**C-D**) pDyn and pEnk probes. (**E**) Illustration of area of PBN depicted **in F-K**. (**F-H**) Representative FISH images of LPBN neurons expressing (**F**) preprodynorphin (pDyn), (**G**) preproenkephalin (pEnk), and (**H**) slc17a6 (VGLUT2). (**I-K**) (similar results were obtained in n=3 mice) Representative imagines of overlays of (**I**) preprodynorphin with slc17a6 and preproenkephalin with slc17a6 (**J**), and (**K**) preprodynorphin with preproenkephalin. Arrowheads mark examples of cells positive for co-labeling of two transcripts. 98% of neurons expressing prodynorphin and 97% of neurons labeled for proenkephalin were also labeled with probes for slc17a6 (VGLUT2). Data are presented as mean ± SEM; n = 4 animals, 760 cells for pDyn and n = 4 animals, 760 cells for pEnk.) Diagram of viral injections into wild-type mice. (**M**) Anatomical location of representative FISH images shown in (**N-O**) that show overlap of **(N**) Cre expression, mediated by retrovirus transduction, with (**O**) preprodynorphin and (**P**) preproenkephalin. Arrowheads mark cells expressing Cre, with filled arrowheads co-expressing (**O**) preprodynorphin or (**P**) preproenkephalin and open arrowheads only expressing Cre.

Next, we examined the connection of pDyn and pEnk expressing neurons in the LPBN to the POA using retrograde AAV’s and FISH. We injected AAV2-retro-Cre into POA of wildtype mice (**Figure 2L**) and collected brain sections containing PBN. We probed these sections for viral induced Cre expression (**Figure 2M**). Using FISH, we observed retrograde viral induced expression of Cre in LPBN (**Figure 2N**) in cells also labeled with pDyn (**Figure 2O**) and pEnk (**Figure 2P**) indicating that neurons expressing these two opioid peptides project to the POA.

To further examine the projections of Dyn+ and Enk+ PBN neurons to the POA, we employed both retrograde AAV’s and anterograde tracing to locate connections from PBN Dyn+ and Enk+ neurons. To identify anterograde projections of PBN neurons, we injected the Dyn-Cre or Enk-Cre mice with AAV5-Ef1a-DIO-eYFP or AAV5-Ef1a-DIO-ChR2-eYFP into the LPBN. To retrogradely label POA projecting neurons we injected AAV2-retro-CAG-FLEX-tdTomato-WPRE into the POA of the same Dyn-Cre or Enk-Cre animals (**Figure 3A, F**). In this, experiment, we observed anterograde labeling of processes with eYFP, from viral injections in the PBN, in the ventral medial preoptic hypothalamus (VMPO) in brains from both Dyn-Cre (**Figure 3C**) and Enk-Cre (**Figure 3H**) mice. Retrograde labeling of LPBN neurons by Cre-dependent expression of tdTomato from retroAVV injected into the POA was evident in sections from both Dyn-Cre (**Figure 3E**) and Enk-Cre (**Figure 3J**) brains. Double-labeled cells, expressing both tdTomato (retrograde) and eYFP were present in the LPBN of both Dyn-Cre and Enk-Cre mice (**arrowheads in Figure 3E and J**). In Sagittal sections of brains taken from Dyn-Cre mice injected with AAV-DIO-ChR2eYFP in the PBN we also observed labeled projections to the POA (**Supplemental Figure 1J-K).**

**Figure 3.**
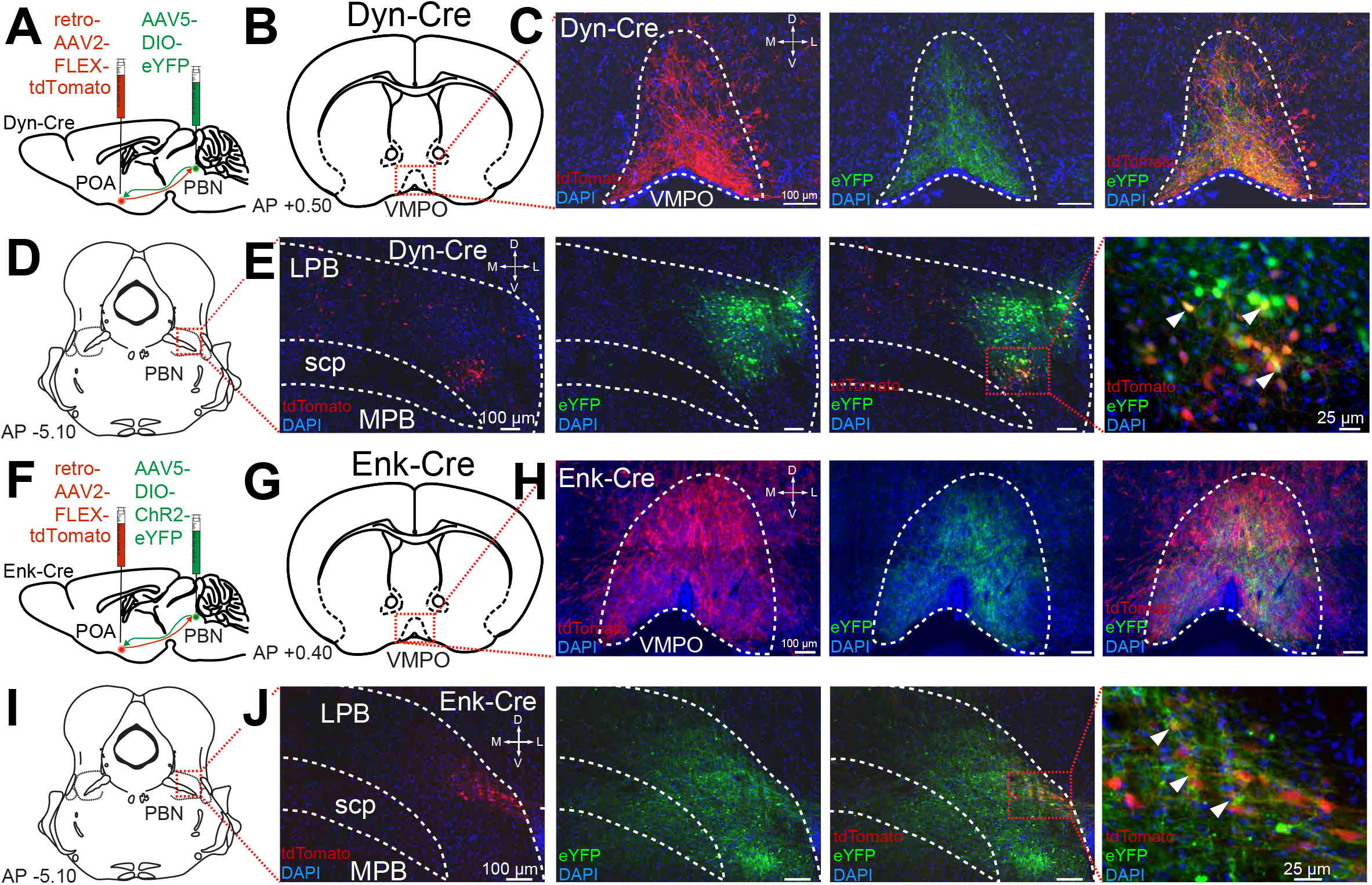
Dyn+ and Enk+ LPBN neurons project to VMPO. (**A**) Illustration of injection of retroAAV-DIO-tdTomato in POA and AAV5-DIO-eYFP in a Dyn-Cre mouse. (**B**) Diagram of POA region depicted in (**C**) showing antero-(green) and retrograde (red) labeling of Dyn+ neurons in POA. (**D**) Diagram of PBN region depicted in (**E**) showing retrograde labeling from POA (red) and eYFP expression (green). Yellow cells in overlay image, marked with arrow heads, illustrate dual labeling by locally injected and retrograde viruses. (**F**) Illustration of injection of retroAAV-DIO-tdTomato in POA and AAV5-DIO-eYFP in an Enk-Cre mouse. (**G**) Diagram of POA region depicted in (**H**) show antero-(green) and retrograde (red) labeling of Enk+ neurons in POA. (**I**) Diagram of PBN region depicted in (**J**) showing retrograde labeling from POA (red) and eYFP expression (green). Yellow cells in overlay image, marked with arrow heads, illustrate dual labeling by locally injected and retrograde viruses.

To examine which neurons comprise the PBN to POA projecting population we injected mice expressing Cre under control of the VGLUT2 (Slc17a6) promoter (VGLUT2-Cre) (Vong et al., 2011) with AAV5-DIO-ChR2eYFP bilaterally in the PBN, labeling VLGUT2 expressing PBN neurons (**Supplemental Figure 1E**). We observed VGLUT2 positive cells labeled by eYFP in the MPBN and LPBN after viral injection (**Supplemental Figure 1F-G**). VGLUT2+ projections from the PBN to the POA including the ventral medial preoptic nucleus (VMPO) and the median preoptic nucleus (MNPO) were labeled by AAV5-DIO-ChR2eYFP injected in the PBN (**Supplemental Figure 1H-I**). To test whether Dyn+ or VGLUT2+ cells represented the whole of the population of PBN to POA neurons, a retrograde recombinase dependent red-to-green (tdTomato to EGFP) color-changing virus (AAV-retro-DO_DIO-tdTomato_EGFP) was injected into the POA of Dyn-Cre or VGLUT2-Cre mice **(Supplemental Figure 1A)**. In Dyn+ Cre mice, we observed cells in LPBN expressing tdTomato (Cre negative cells) and neurons expressing eGFP (Cre positive cells) (**Supplemental Figure 1C)**. In VGLUT2+ mice, we only observed eGFP expressing (Cre positive cells) neurons in LBPN (**Supplemental Figure 1D)** indicting the PBN to POA projection is composed entirely of VGLUT2+ cells. Taken together, results from FISH experiments and viral tracing studies indicate that Dyn+ and Enk+ neurons in LPBN project to the POA, particularly the VMPO, and that both Dyn+ and Enk+ POA projecting neurons are subsets of the VGLUT2+ population of LPBN neurons that project to POA.

### Photostimulation of Dyn^PBN→POA^, Enk^PBN→POA^, and VGLUT2^PBN→POA^ generates rapid onset of hypothermia

Using the respective Cre driver lines, we next examined the roles of POA-projecting Dyn+, Enk+, and VGLUT2+ PBN neurons (circuits are denoted as Dyn^PBN→POA^, Enk^PBN→POA^, and VGLUT2^PBN→POA^, respectively) in regulating body temperature. We injected AAV5-DIO-ChR2-eYFP bilaterally into the LPBN of Dyn-Cre, Enk-Cre, and VGLUT2-Cre mice, and after 6-weeks, we implanted a single midline optic fiber above VMPO, where projections from PBN were observed (**Figure 4A-B**). We implanted mice with a subdermal wireless temperature transponder to enable touch free recording of body temperature. For each trial, we connected mice to an optic patch cable, and following a one-hour period of habituation to the behavioral arena, we photostimulated PBN→VMPO terminals for 15 minutes with 10 ms light pluses at pulse frequencies of 2, 5, 10, and 15 Hz (**Figure 4C**). We recorded body temperature every 5 minutes for 65 minutes, beginning 5 minutes prior to photostimulation (**Figure 4C**).

**Figure 4.**
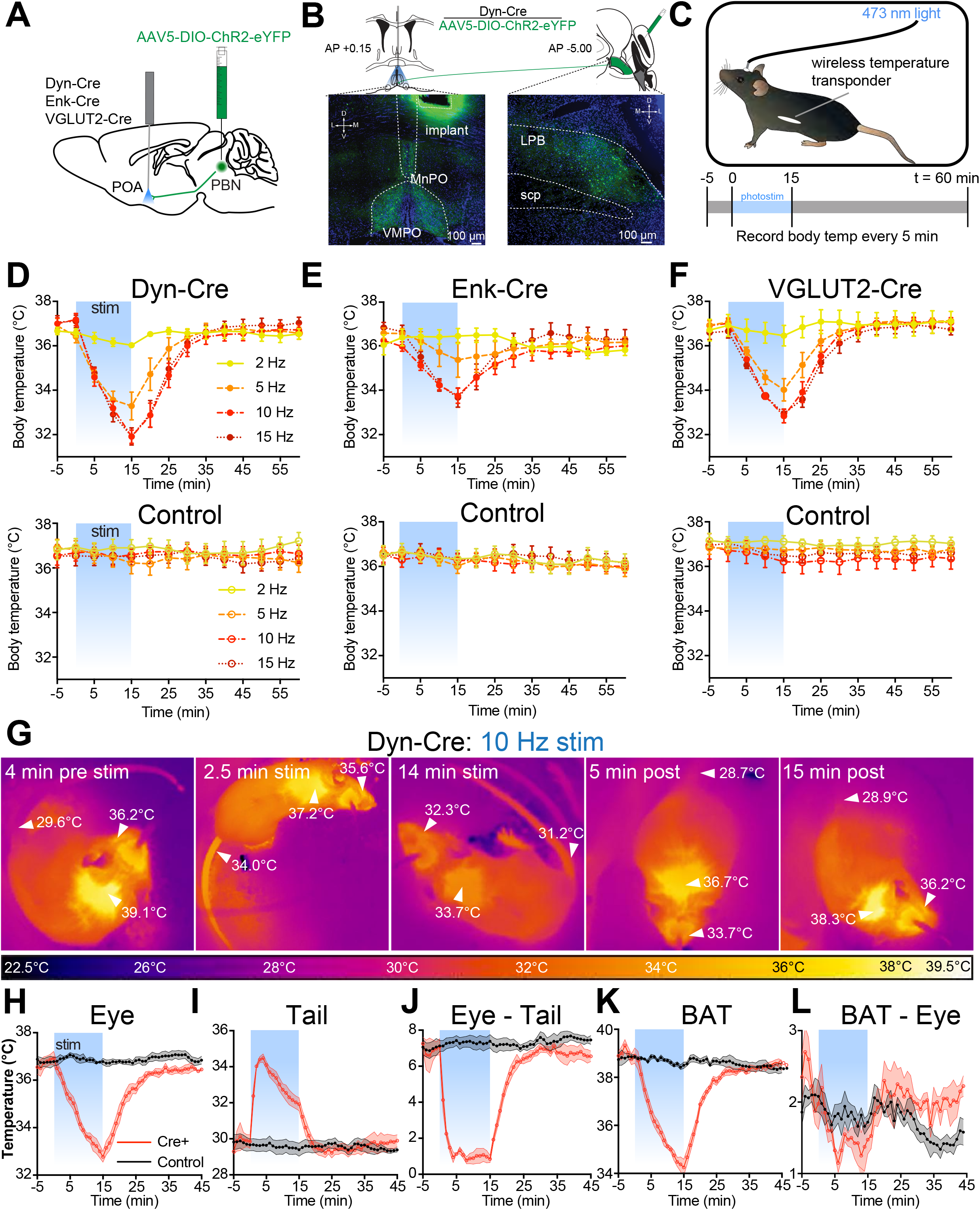
Photostimulation of Dyn^PBN→POA^, Enk^PBN→POA^, and VGLUT2^PBN→POA^ leads to acute hypothermia by evoking thermal heat defenses. (**A**) Illustration of viral injections in PBN and fiber optic implantation over POA in Dyn-Cre, Enk-Cre, or VGLUT2-Cre mice. (**B**) Illustration shows viral and fiber optic delivery in a Dyn-Cre mouse along with representative expression of ChR2-eYFP (green) in PBN injection site and POA implantation site. (**C**) Diagram shows core body temperature measurement method and paradigm for photostimulation for 15 min and temperature recording for 65 min trials. (**D-F**) Body temperature vs. time graphs for 2 (yellow), 5 (orange), 10 (red), and 15 (dark red) Hz photostimulation of (**D**) Dyn^PBN**→**POA^, (**E**) Enk^PBN**→**POA^, (**F**) VGLUT2^PBN**→**POA^, and controls for each. Photostimulation was delivered from t=0 to t=15 min and led to a frequency dependent reduction in body temperature in Dyn-Cre, Enk-Cre, and VGLUT2-Cre mice. Body temperature of control animals was stable throughout the trials. Data are presented as mean ± SEM. For experimental animals, n=6 (**D and E**) and n=8 (**F**). For control animals, n=8 (D) and n=7 (**E and F**). (**G**) Representative quantitative thermal imaging from a representative trial showing a mouse before, during, and after 10 Hz photostimulation of Dyn^PBN**→**POA^. Arrows show temperatures of eye, BAT, or tail. Eye and BAT temperature decreased as a result of stimulation; tail temperature increased as a result of stimulation. (**H-L**) Quantitative thermal imaging measurements of (H) eye, (**I**) tail, (**J**) Eye minus Tail, (**K**) BAT, and (**L**) BAT minus Eye temperature vs. time graphs for 10 Hz photostimulation of Dyn^PBN**→**POA^. Photostimulation was delivered from t=0 to t=15 min and led to decreases in eye and BAT temperatures, an increase in tail temperature. Tail and eye temperatures equilibrated in Cre+ animals. BAT thermogenesis was suppressed with a decline in the difference between eye and BAT temperatures during stimulation. Data are presented as mean ± SEM. See Supplemental Figure 2 for data from Enk-Cre animals.

Photostimulation of Dyn^PBN→POA^ neuron terminals caused rapid and significant reduction in body temperature in Dyn-Cre mice (n=6), with increasing magnitude of drop in body temperature corresponding to increasing photo-stimulation frequency up to 10Hz (**Figure 4D**). 15 mins of stimulation of Dyn^PBN→POA^ projections reduced the body temperature to 36.0±0.1°C at 2 Hz (p=0.571 vs. control), 33.3±0.6°C (p=.0032) at 5Hz, 31.9±0.3°C, (p<0.0001) at 10Hz, and 31.9±0.4°C (p<0.0001) at 15Hz compared to control. In control mice (n=7), photostimulation did not cause significant changes in body temperature at any of the tested frequencies (**Figure 4D**).

Photostimulation of Enk^PBN→POA^ neuron terminals also caused a rapid reduction in body temperature (**Figure 4E**) in a stimulation frequency dependent manner. 15 mins of stimulation of Enk^PBN→POA^ projections in Enk-Cre mice reduced body temperature to 36.4±0.3°C at 2 Hz (p=0.999 vs. control), 34.9±0.7°C (p=0.495) at 5 Hz, 33.8±0.3°C (p=0.0001) at 10Hz, and 33.7±0.4°C (p=0.0002) at 15 Hz, compared to a separate cohort of control mice (n=7) which did not display altered body temperatures in response to photo stimulation.

In VGLUT2-Cre mice with AAV-DIO-ChR2eYFP injected into PBN, stimulation of VGLUT2^PBN→POA^ terminals in POA also caused a rapid and significant decrease in body temperature (**Figure 4F**). 15 minutes of photostimulation in VGLUT2-Cre mice (n=8) significantly reduced the mean ±SEM body temperature to 36.5 ±0.5°C at 2Hz (p=0.257 vs. control) 34.0 ±0.5°C at 5Hz (p=0.0005), 32.8 ±0.3°C (p<0.0001) at 10Hz, and 32.9 ±0.2°C (p<0.0001) at 15Hz compared to control mice (n=7). The average changes in body temperature that we measured in Dyn-Cre and VGLUT2-Cre mice were not significantly different at any of the tested stimulation frequencies. The body temperature reduction evoked by photostimulation in Enk-Cre mice was smaller in magnitude than that in either Dyn-Cre or VGLUT-Cre mice. The mean body temperature we measured in Enk-Cre mice after 15 minutes of simulation was significantly different than Dyn-Cre at 10 Hz (p=0.02), with activation of the Enk+ terminals having less of an effect. These data demonstrate that activation of PBN→POA terminals causes rapid decreases in body temperature.

### Photostimulation of Dyn^PBN→POA^ and Enk^PBN→POA^ terminals causes vasodilation and suppresses brown fat thermogenesis

We sought to examine mechanisms causing core body temperature reduction in response to photostimulation of PBN→POA projections. We used thermal imaging to measure temperatures of eye, tail, and interscapular region, which covers brown adipose tissue (BAT), in Dyn-Cre mice (representative imaging in **Figure 4G**). Thermal imaging of the eye has previously been demonstrated as an accurate proxy for core body temperature (Vogel et al., 2016). We recorded eye temperatures every minute during a 10Hz photostimulation paradigm, as described above. Recorded eye temperatures demonstrated a rapid reversible decrease after photostimulation (**Figure 4H**) and closely tracked values obtained using implanted wireless transponders. In Dyn-Cre mice, mean ±SEM eye temperature dropped from 36.9°C ±0.3 to 32.8°C ±0.2 with 15 mins of stimulation (**Figure 4H**). Thermal imaging to quantify tail temperature can be used to observe heat loss from vasodilation in response to warmth (Meyer et al., 2017). We obtained thermal imaging measurements of the tail temperatures approximately 1 cm from the base of the tail each minute. In Dyn-Cre mice, tail temperature measurements demonstrated a very rapid increase following the onset of photostimulation, increasing a mean ±SEM of 4.2°C ±0.5 after 2 minutes of photostimulation **(Figure 4I**). Increase in tail temperature preceded the decline in core body temperature. As core body temperature began to decrease, the tail temperature also began to decrease (**Figure 4H and I**). We examined the difference between the tail and eye temperatures (**Figure 4H-J**) to determine whether the gradient between core and peripheral temperature was maintained as body temperatures declined during stimulation. At baseline we observed a mean ±SEM difference 6.9 ±0.29°C between the measured eye and tail temperatures. Eye-tail temperature difference significantly (p<0.0001) decreased compared to control to a mean ±SEM of 1.3 ± 0.3 °C and remained stable during photo stimulation even as body temperature declined. The difference in eye-tail temperature returned to baseline shortly after photostimulation was stopped **(Figure 4J**).

Previous studies have implicated the POA in regulating BAT metabolism and temperature in response to cooling (Nakamura and Morrison, 2007; Tan et al., 2016). To simultaneously examine the response of BAT to the PBN→POA photostimulation-induced hypothermia, temperature measurements were also made of the interscapular BAT region temperature in mice with the fur removed from over the intrascapular region. In Dyn-Cre mice, the temperature of the BAT region decreased rapidly following the onset of stimulation and returned to baseline post-stimulation in a pattern similar to body temperature (**Figure 4K**). If BAT activity responded to the decrease in body temperature by increasing metabolism, then the BAT-eye temperature difference would be expected to increase, reflecting the warming activity of BAT and the falling body temperature. The temperature difference between BAT and eye (BAT-Eye) decreased during the period of stimulation but returned to baseline when stimulation was stopped (**Figure 4L**). We conducted similar experiments using thermal imaging in Enk-Cre and additional control animals (**Supplemental Figure 3**). In the Enk-Cre mice, we found similar effects, but of smaller overall amplitude. Photostimulation of Enk^PBN→POA^ terminals led to a decrease in eye temperature, a rapid increase in tail temperature, decrease in BAT temperature, and collapse of the eye-tail temperature gradient (**Supplemental Figure 3D-G**). Together these results indicate that PBN to POA neurons can drive physiologic adaptation to lower body temperature by increasing heat dissipation and suppressing thermogenesis.

### Changes in body temperature evoked by photostimulation of Dyn^PBN→POA^ and Enk^PBN→POA^ terminals are opioid peptide and receptor independent

To test the potential role of endogenous opioids and their receptors in mediating the alterations in body temperature evoked by activation of Dyn^PBN→POA^ and Enk^PBN→POA^ terminals, mice were treated with opioid receptor antagonists prior to photostimulation (**Supplemental Figure 3A)**. Dyn-Cre (n=7) and control mice (n=7) were treated with the opioid receptor antagonist naltrexone (3mg/kg) via intraperitoneal (IP) injection and then given a 10Hz photostimulation paradigm as above (**Supplemental Figure 3B**). The order of naltrexone and saline was varied between animals, and trials were run on separate days. 30 minutes after treatment with naltrexone, we did not observe a significant impact on photo-stimulation induced change in body temperature compared to saline treated animals. Naltrexone was paired with the Dyn-Cre line because of the relatively higher affinity of naltrexone for kappa opioid receptors compared to naloxone (Meng et al., 1993). A distinct cohort of Dyn-Cre mice (n=4) was treated with saline and for subsequent trials with Norbinaltorphimine (NorBNI) 10 mg/kg via IP injection 1 day prior and again 30 minutes prior to photostimulation. Pretreatment with NorBNI did not significantly alter the decrease in body temperature induced by 10Hz photostimulation of Dyn^PBN→POA^ terminals (**Supplemental Figure 3B**). Enk-Cre mice (n=5) injected with AAV5-DIO-ChR2eYFP in the PBN and control mice (n=4) were treated with naloxone 5 mg/kg and saline, with the order of treatments varied between animals and trials conducted on separate days. 30 minutes after treatment with naloxone, Enk^PBN→POA^ terminals were photostimulated at 10 Hz. No significant effect of naloxone on photostimulation-induced changes in body temperature was observed (**Supplemental Figure 3C)**. These data suggest that the acute alterations in body temperature due to stimulation of Dyn^PBN→POA^ and Enk^PBN→POA^ terminals are not driven by endogenous opioid release and subsequent opioid receptor signaling.

### Glutamatergic PBN neuronal activity is necessary for heat-induced vasodilation

Glutamatergic signaling in the POA and in PBN has previously been implicated in heat defensive behaviors (Nakamura and Morrison, 2010), and our results, presented here, demonstrate sufficiency of PBN VGLUT2+ neurons in driving hypothermia (**Figure 4F**). To examine the necessity of VGLUT2+ PBN neurons in mediating heat defensive behaviors in awake behaving animals, AAVs encoding Gi coupled DREADDs in a Cre-dependent manner (AAV-DIO-hM4DGi) were injected into PBN bilaterally in VGLUT2-Cre mice (n=5). Mice were treated with saline or clozapine-N-oxide (CNO) (2.5 mg/kg IP) 30 minutes prior to a heat challenge of 34°C for 15 minutes and were tested with the reciprocal during a subsequent trial more than 24 hours later. Here we used a custom small arena with floor and walls lined with a water jacket connected to circulating water baths at 20°C or 34°C to create a rapid change in temperature between two stable set points while allowing for continuous thermal imaging (**Figure 5A–B)**. Using quantitative thermal imaging, we measured tail temperatures and arena floor temperatures (depicted by the yellow-orange shaded areas) every minute during chemogenetic inhibition of VGLUT2 activity (**Figure 5B-D**). In mice treated with saline, measurements of tail temperatures showed a rapid rise following the shift of arena temperature to 34°C and measured tail temperatures were higher than the arena floor temperature (**Figure 5B, D and F**). In mice treated with CNO, which activated the inhibitory DREADD in VGLUT2+ PBN neurons, the mean ±SEM tail temperature after 15 minutes of exposure to 34°C was 34.8°C ±0.5, significantly (p=0.01) lower than the corresponding average tail temperature after saline treatment, 37.1°C ±0.5 (**Figure 5D**). The tail temperature in saline treated mice exceeded the temperature of the arena floor **(Figure 5B and F**), but in CNO treated mice, tail temperature rose only to the temperature of the floor (**Figure 5B and E**). Consistent with an effect of passive heating of the tail, as opposed to the active vasodilation evoked by the thermal challenge, the rate of increase in the tail temperature was also slower following CNO treatment compared to saline (**Figure 5C**). After return of the arena floor temperature to 20°C, tail temperatures returned to a baseline of approximately 22°C following both saline and CNO treatments. CNO treatment in WT mice had no significant effects on tail temperature changes compared to saline treatment (**Supplemental Figure 4).** These results indicate that VGLUT2+ PBN neurons are required for heat defensive responses including physiological vasodilation.

**Figure 5.**
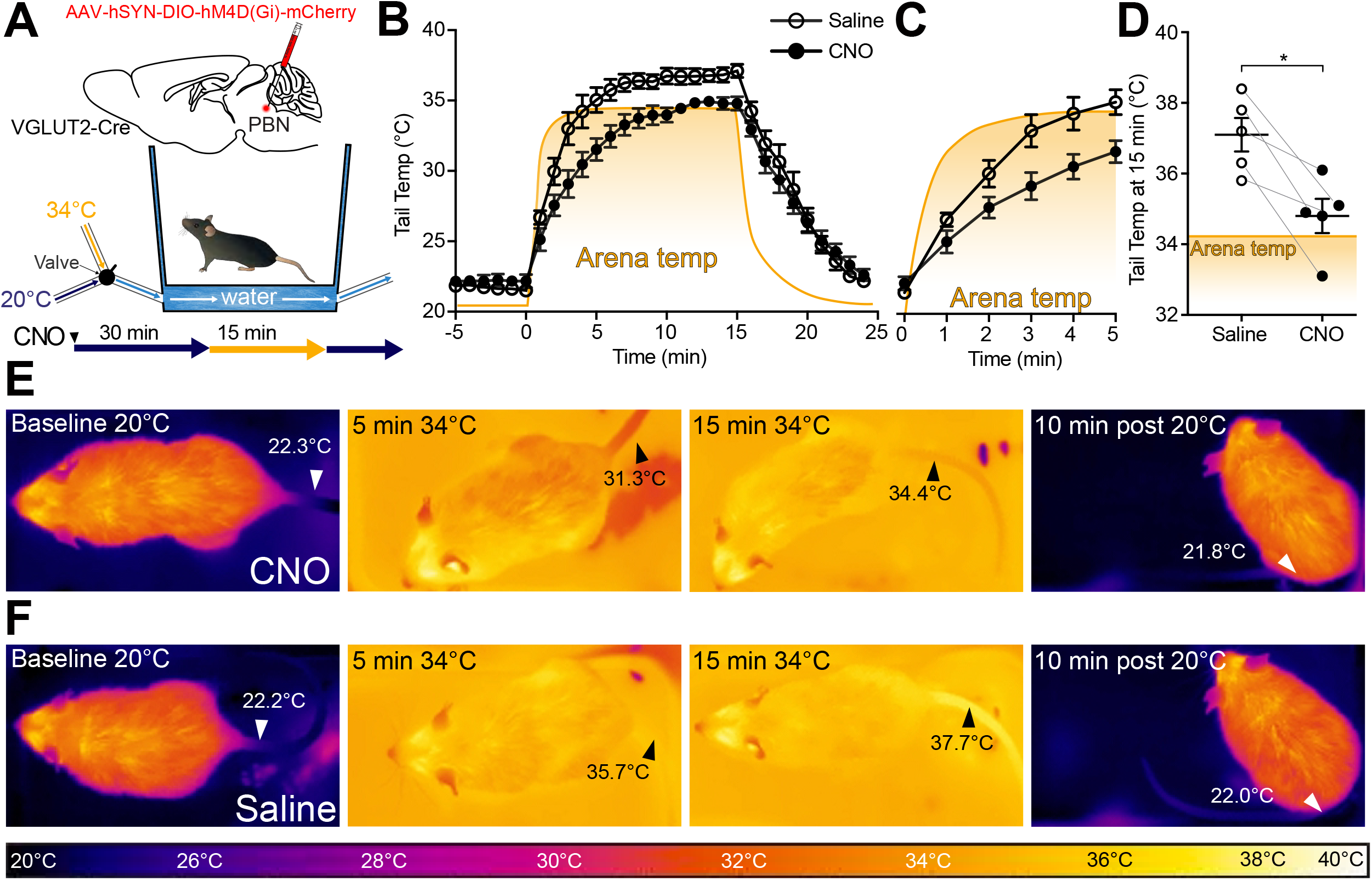
VGLUT2+ PBN neurons are necessary for heat-defensive tail vasodilation. (**A**) Illustrations depict viral injections in VGLUT2-Cre mice and purpose-built heat challenge arena that allowed for rapid changing of environmental temperature between two stable set points. (**B**) Tail temperature as determined using quantitative thermal imaging vs. time graph for 34°C thermal heat challenge for mice expressing hM4D(Gi) DREADDs in VGLUT2+ PBN neurons treated either with CNO or saline. Heat challenge was delivered from t=0 to t=15 min, and arena temperature measured using thermal imaging during the trial is represented by the orange line. In mice injected with CNO 2.5 mg/kg, tail temperature passively equilibrated with arena temperature (34°C) over the 15-minute heat challenge. In mice injected with saline, tail temperature rose above arena temperature after 5 minutes of heat challenge representing heat release through vasodilation. Data are presented as mean ± SEM. n=5 animals, paired between CNO and saline conditions.(**C**) Tail temperature vs. time graph for 34°C heat challenge between t=0 and t=5 min. Note the separation between average tail temperatures of the saline condition vs. the CNO condition. Data are presented as mean ± SEM. n=5 animals, paired between CNO and saline conditions. (**D**) Tail temperature at t=15 min of 34°C heat challenge. Tail temperatures in the saline condition were an average of 2.3±0.68°C higher than those in the CNO condition. (**E**) Representative thermal images of trials for mice treated with CNO and measurement of tail temperature showing tail temperatures remain close to the temperature of the area floor. (**F**) Representative thermal images of trials for mice treated with saline and tail temperature exceed floor temperature. Data are presented as mean ± SEM. n=5 animals, paired between CNO and saline conditions. Student’s t test, *p < 0.05. See Supplemental Figure 3 for data from the same assay in Dyn-Cre mice.

### Photostimulation of PBN→POA drives thermal defensive behaviors

To test the sufficiency of Dyn^PBN**→**POA^, Enk^PBN**→**POA^, and VGLUT2^PBN**→**POA^ to cause specific avoidance behavior we conducted real-time place aversion (RTPA) experiments using the respective Cre driver lines and photostimulation of terminals in the POA. Photostimulation of terminals was paired to entry into one compartment of a balanced two-compartment conditioning apparatus void of salient stimuli. Neurons that encode a negative valence will cause an aversion from the chamber paired with photostimulation, and those with a positive valence will drive a preference for it (Kim et al., 2013; Namburi et al., 2016; Siuda et al., 2015; Stamatakis and Stuber, 2012; Tan et al., 2012). As in experiments above, we injected AAV-DIO-ChR2-eYFP into the PBN bilaterally of Cre driver line mice and implanted optical fibers over the POA (**Figure 6A, D, G**). Photostimulation of Dyn^PBN**→**POA^ terminals drove aversion in a frequency dependent manner, with time spent in the stimulation side being significantly lower (p<0.0001) at 5, 10, and 20 Hz stimulation frequencies compared to control mice (**Figure 6A-C)**. Results from parallel RTPA experiments using Enk-Cre mice demonstrated a similar effect of aversion seen at 5 (p=0.0002), 10 (p<0.0001), and 20 Hz (p<0.0001) stimulation frequencies compared to control mice (**Figure 6D-F**). Results we obtained using VGLUT2-Cre mice in RTPA experiments showed significant (p<0.0001) decreases in time spent on the stimulation side at 2, 5, 10, and 20 Hz compared to control animals (**Figure 6G-I**). For each genetic line, we examined the locomotor activity or distance traveled during the RTPA. In Dyn-Cre mice, we observed a small but significant (p=0.009) difference in mean ±SEM total distance traveled only during trials using 20Hz stimulation - 29m ±2 (n=8) compared to control 48m ±5 (n=7) - but not during trials using lower stimulation frequencies **(Supplemental Figure 5A**). In Enk-Cre and VGLUT2-Cre mice, we observed no significant differences between Cre+ and control animals at any of the photo-stimulation frequencies (**Supplemental Figure 5B, C**). Comparisons of male vs female mice did not reveal sex dependent effects in the acute hypothermic changes in body temperature evoked by photostimulation of PBN→POA terminals (**Supplemental Figure 5D-E**).

**Figure 6.**
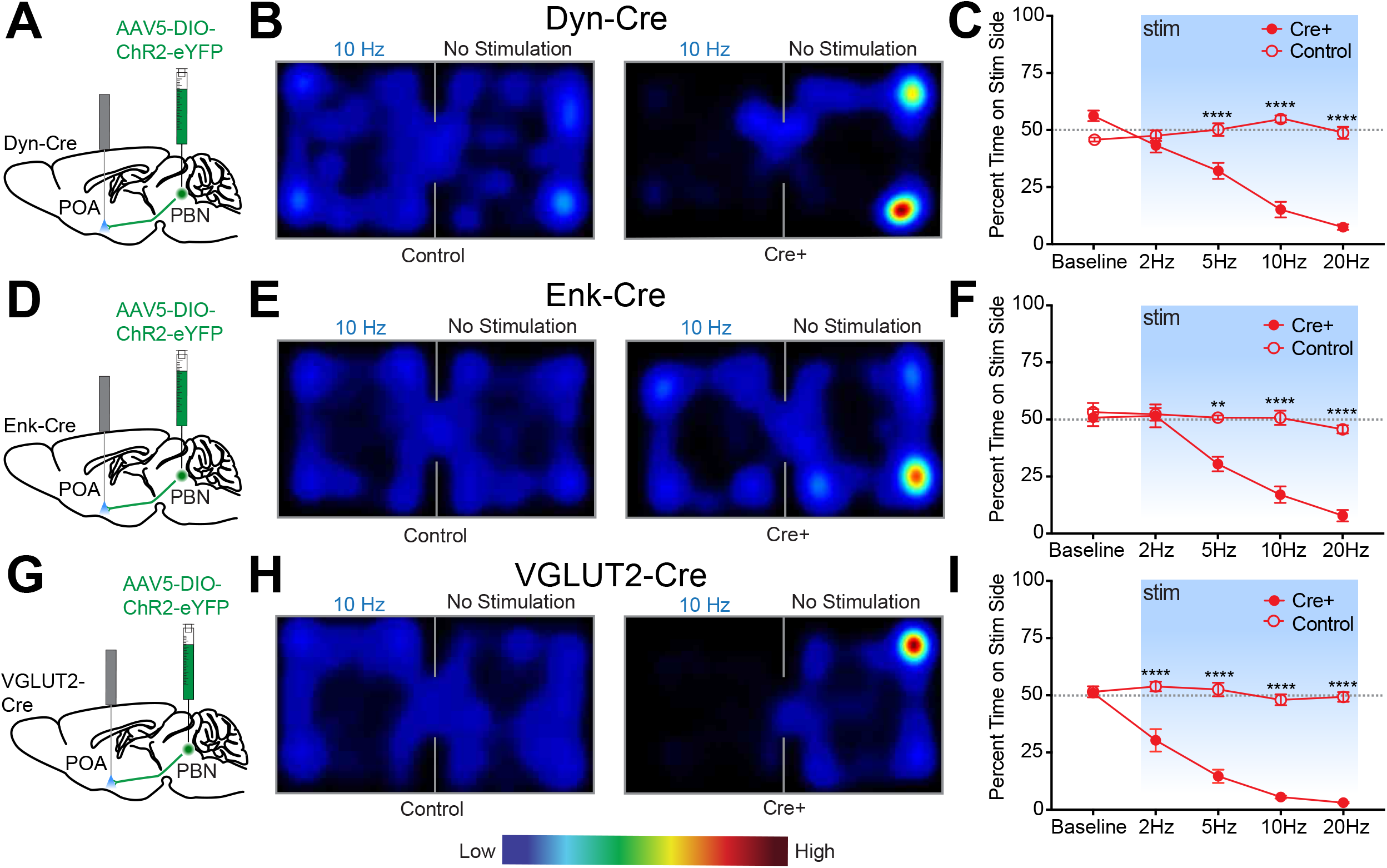
Photostimulation of Dyn^PBN→POA^, Enk^PBN→POA^ and VGLUT2^PBN→POA^ terminals drives real time place aversion. (**A, D, and G**) Illustrations of viral injections in PBN and fiber optic implantations over POA in Dyn-Cre mice, Enk-Cre and VGLUT2-Cre mice, respectively. (**B, E and H**) Representative heat maps showing spatial distribution of time-spent behavior resulting from side-conditional 10 Hz photostimulation of control or Dyn-Cre, Enk-Cre and VGLUT2-Cre mice, respectively. (**C**) For Dyn-Cre vs control mice, frequency response of RTPP at 0 (baseline), 2, 5, 10, and 20 Hz. Data are presented as mean ± SEM; n=6 Cre+, 8 control; two-Way ANOVA, Bonferroni post hoc ((**F**) Enk-Cre frequency response of RTPP at 0 (baseline), 2, 5, 10, and 20 Hz. Data are presented as mean ± SEM; n=6 Cre+, 7 control; two-Way ANOVA, Bonferroni post hoc (5 Hz ChR2 vs. 5 Hz control ***p < 0.001, 10 Hz ChR2 vs. 10 Hz control ****p < 0.0001, 20 Hz ChR2 vs. 20 Hz control ****p < 0.0001) (**I**) VGLUT2-cre frequency response of RTPP at 0 (baseline), 2, 5, 10, and 20 Hz. Data are presented as mean ± SEM; n=8 Cre+, 7 control; two-Way ANOVA, Bonferroni post hoc (2 Hz ChR2 vs. 20 Hz control ****p < 0.0001, 5 Hz ChR2 vs. 5 Hz control ****p < 0.0001, 10 Hz ChR2 vs. 10 Hz control ****p < 0.0001, 20 Hz ChR2 vs. 20 Hz control ****p < 0.0001). See also Supplemental Figure 4.

Other thermoregulatory behaviors including posture, stance, and locomotion are altered by exposure to warm temperatures (Cabanac, 1975). Therefore, we next examined alterations in locomotion using 20-minute open field-testing trials in Dyn-Cre and control mice (**Figure 7A-C**). Stimulation of Dyn^PBN→POA^ terminals at 10Hz resulted in a large and significant (p=0.0008) decrease in mean ±SEM distance traveled: 26.1m ±6.2 (n=5) in Dyn-Cre compared to 65.9m ±5.6 in control mice (n=7) (**Figure 7B-C**). Postural extension, depicted in the photograph (**Figure 7D**), a heat evoked behavior in rodents that reduces heat production by postural tone and increases exposed body surface to promote thermal transfer (Roberts, 1988), was evoked by photostimulation of Dyn^PBN→POA^ terminals. Scoring of video recordings of trials of Dyn-Cre (n= 7) and control mice (n=4) revealed that Dyn-Cre mice quickly transition to a sprawled posture after the onset of 10 Hz photostimulation, which also induces hypothermia. Following the end of photostimulation, mice transition and spend more time in a posture with their tail curled under their bodies to minimize exposed surface area (**Figure 7E**). We did not observe postural extension at any time in control mice during these trials.

**Figure 7.**
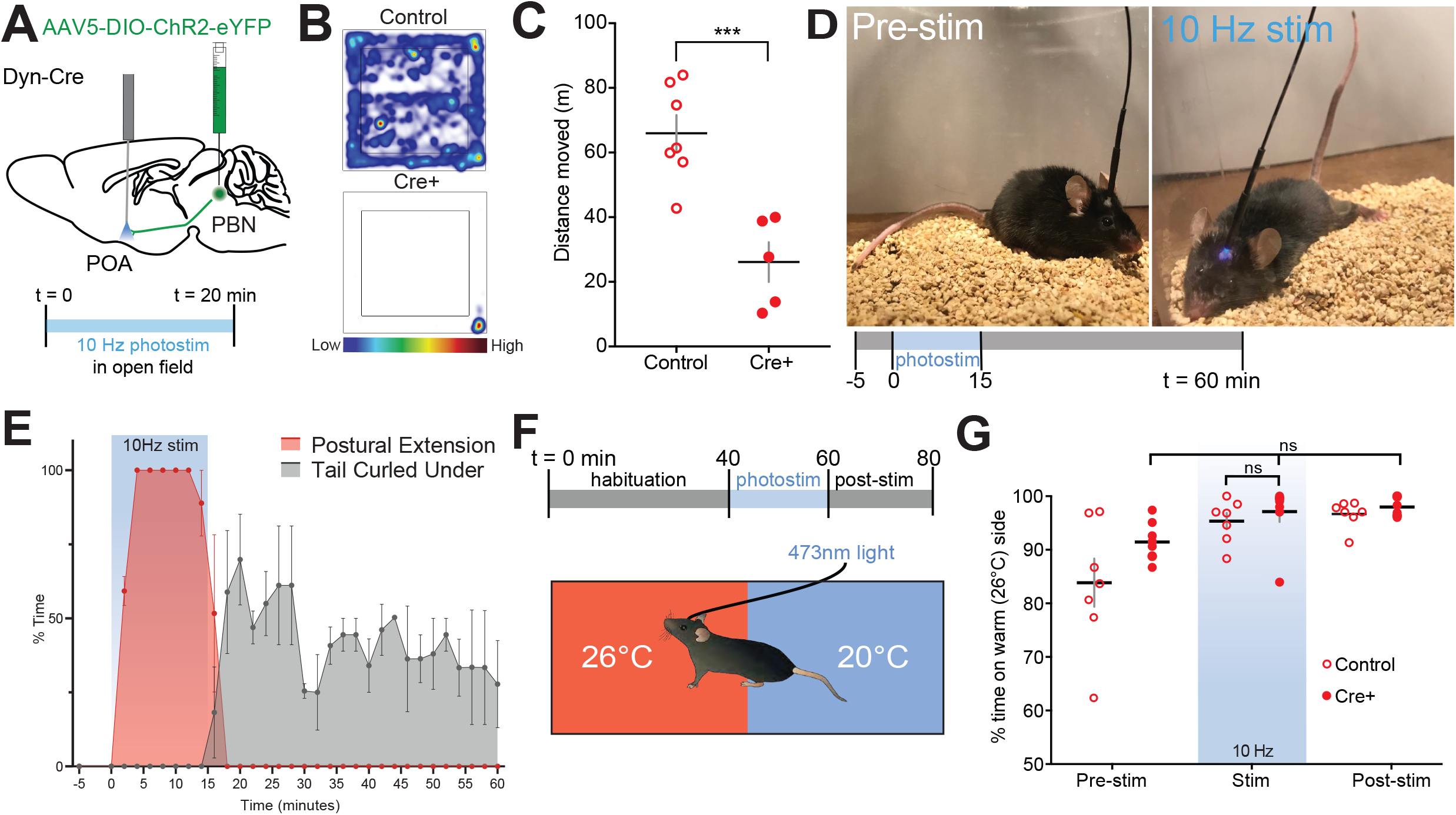
Photostimulation of Dyn^PBN→POA^ suppresses locomotion, evokes postural extension but does not alter temperature preference. (**A**) Illustration of injection in PBN and fiber implantation over POA in Dyn-Cre mice. (**B**) Representative heat maps show spatial distribution of time-spent behavior resulting from constant 20 min 10 Hz photostimulation of control or Dyn^PBN**→**POA^. (**C**) Quantification of movement during open field testing. Control animals moved an average of 39.84±8.33 meters more than Cre+ animals during open field trials. Data are presented as mean ± SEM; n=5 Cre+, 7 control; Student’s t test, ***p < 0.001 (**D**) 10 Hz photostimulation of Dyn^PBN**→**POA^ leads to postural extension behavior as shown. Representative images of a mouse pre stimulation and during 10 Hz photostimulation of Dyn^PBN**→**POA^. (**E**) Quantification of percent time spent in time spent engaged in postural extension in Dyn-Cre mice in two min time bins. Following onset of photostimulation mice engaged in postural extension (red). With termination of stimulation mice, we noted to switch to a posture with their tails curled under their bodies (grey). Postural extension was not observed in any control mice. (**F**) Overview of paradigm with 3 epochs: 40 minutes of pre-stim, 10 Hz photostimulation for 20 minutes, and post-stim for 20 minutes in an arena with aluminum floor held at 20°C and 26°C on opposing sides. (**G**) Quantification of time spent in each temperature area showed non-significant changes in percent time spent in each area during delivery of stimulation, with a strong preference for the 26°C side during all epochs. Data presented as mean ±SEM with individual values, n=9 Dyn-Cre (ANOVA ns=0.7341 for Dyn-Cre mice across epochs) and (t-test ns p>.99 for Dyn-Cre vs Control during stimulation epoch).

Temperature selection is an important complex thermal defense behavior. Moving to an area with cooler environmental temperature, when possible, is a way to defend against excessive heat. Available studies indicate thermal selection requires the engagement of multiple poorly understood neural circuits. We next tested whether photoactivation of Dyn^PBN→POA^ terminals is sufficient to induce a shift to slightly cooler temperature preference. To examine temperature preference, we placed mice in an arena with an aluminum floor in which each side is held at a set temperature of 20°C or 26°C (**Figure 7F**). Mice were habituated to the arena prior to the start of the trial to familiarize the animals to area. Trials consisted of a 40 min pre stimulation period, a 20 min stimulation period, and a 20 min post stimulation period. As expected, at baseline, mice spent a greater amount of time on the 26°C side (**Figure 7G)**. In Dyn-Cre mice (n=6), photostimulation of the POA at 10 Hz did not alter animals’ thermal preference to the cooler side of arena (**Figure 7**), despite this photostimulation paradigm evoking hypothermia (**Figure 4**) and driving other thermal defense behaviors. This result indicates that the Dyn^PBN→POA^ neurons are not sufficient to drive cool seeking behavior when activated, suggesting that other neural pathways are also required to drive this behavior.

## Discussion

In the present study we demonstrate that warm-activated neurons within the PBN overlap with neural populations (Dyn+ and Enk+) marked by Cre reporters for expression of *preprodynorphin* and *preproenkephalin* (**Figure 1 and Supplemental Figure 2**). Employing FISH, we found that PDyn+ and PEnk+ expressing neuronal populations also express VGLUT2, partially overlap, and project to the POA (**Figure 2**). Using anterograde and retrograde viral tools, we demonstrate that Dyn+, Enk+, and VGLUT2+ PBN neurons project to the POA (**Figure 3 and Supplemental Figure 1)**. We found that photoactivation of Dyn+, Enk+ and VGLUT2+ PBN terminals in POA is sufficient to drive physiological (**Figure 4**) and behavioral heat defense behaviors (**Figures 6, 7**).

### Overlapping populations of warm-activated PBN neurons express Enk and Dyn, are glutamatergic, and project to the POA

Here we report that a large percentage of PBN neurons are activated following exposure to warmth. The majority of warm-activated PBN neurons express Dyn+ and, surprisingly, a smaller population of these warm-activated PBN cells are Enk+ (**Figure 1**). The 81% and 54% of cFos+ cells that are Dyn+ and Enk+, respectively, suggests that Dyn+ and Enk+ populations overlap, as The sum of the of cFos+ cells that are Dyn+ and Enk+, 81% and 54% exceeds respectively, 100% Dyn+ and Enk+ populations overlap. FISH experiments *preprodynorphin* and *preproenkephalin* showed that 51% of *preprodynorphin* positive cells were also positive for *preproenkephalin* and that 58% of *preproenkephalin* positive cells were also positive for preprodynorphin (**Figure 2**). Fluorescent *in situ* hybridization results also indicate that preprodynorphin and preproenkephalin expressing cells are glutamatergic based on co-expression of VGLUT2 (**Figure 2**). These findings are consistent with previous reports that implicated glutamatergic FoxP2+ and Dyn+ neurons in the dorsal lateral PBN in responding to warmth (Geerling et al., 2016).

The warm-activated neuron population in the PBN overlaps with Dyn+ and Enk+ populations that project to the POA and is glutamatergic (VGLUT2+). We found dense Dyn+ and Enk+ projections from the PBN to POA including the VMPO (**Figure 3 and Supplemental Figure 1K**). Dyn+, Enk+ and VGLUT2+ PBN cell bodies were labeled by retroAAVs injected into the POA (**Figure 3, Supplemental Figure 1**). Cre dependent color changing, red (tdtomato) to green (GFP), retroAAV injections into the POA indicate that all PBN to POA projecting cells are VGLUT2+ while Dyn+ cells are a subset of the total PBN to POA population (**Supplemental Figure 1A-D**). Cells positive for c-Fos after warmth exposure were subpopulations of the total Dyn+ and Enk+ cells in the PBN. The PBN projects to multiple areas in the POA, such as the VLPO. The PBN also regulates other homeostatic processes including water balance and arousal (Gizowski and Bourque, 2018; Qiu et al., 2016), and PBN to POA projections may be important in an array of homeostatic process. VGLUT2+ projections are seen in POA in the studies here (**Supplemental Figure 1**). Within the PBN, anatomical segregation of functionally discrete circuit pathways is likely important. For example, cold responsive neurons have also been described in the PBN, and cold and warm responsive neurons are anatomically separated, with warm responsive neurons located more caudally in the PBN (Geerling et al., 2016). The projection targets of these PBN cold-responsive neurons remains to be determined in future efforts.

Likely postsynaptic targets in POA for warm-activated PBN cells include the recently identified warmth activated neurons in the POA that express the neuropeptides brain-derived neurotrophic factor (BDNF) and pituitary adenylate cyclase-activating polypeptide (PACAP) (Tan et al., 2016). Also, prior studies have shown that hM3-Gq-DREADD induced stimulation of glutamatergic VMPO neurons (expressing the receptor for the hormone leptin) causes a reduction in core body temperature similar in magnitude to the effect seen by activation of Dyn^PBN→POA^ terminals we observed in the present study. Activation of leptin receptor expressing VMPO neurons also causes mice to display similar postural extension behavior as we observed following activation of Dyn^PBN→POA^ terminals (**Figure 7**) (Yu et al., 2016). Furthermore, glutamatergic neurons in MnPO, which can drive vasodilation, are potential likely targets of the PBN warm-activated cells (Abbott and Saper, 2018). Expression of KOR or MOR in these POA neuronal populations has not been delineated, but injections of opioid receptor agonists into the POA has been shown to alter body temperature indicating that opioid receptors, either pre or post synaptic to the PBN terminals, may have important functional neuromodulatory roles in thermoregulation (Xin et al., 1997). In fact, recently POA KOR signaling was found to modulate body temperature and weight loss during calorie restriction (Cintron-Colon et al., 2019) and deletion of the KOR gene alters weight gain induced by a high fat diet by modulating metabolism (Czyzyk et al., 2010). Opioid receptors are expressed in the PBN, and a presynaptic site of action on PBN terminals may also be relevant to thermal regulation (Chamberlin et al., 1999; De Oliveira et al., 2008; Mansour et al., 1994; Palmiter, 2018; Wolinsky et al., 1996). Future studies using mouse transgenic lines with floxed opioid receptor genes will be useful in determining where within thermoregulatory circuits opioid receptor signaling is critical. Studies are also required to identify the post synaptic targets of PBN neurons in POA and how these neurons integrate information from multiple pathways to regulate thermal homeostasis.

### PBN to POA projections regulate body temperature by evoking physiological and behavioral responses

Here we report results obtained in awake freely behaving mice that demonstrate the role of PBN to POA projecting neurons in driving physiological and behavioral responses to warm thermal challenge. Selective photostimulation of PBN→POA terminals in the three Cre lines (Dyn, Enk, and VGLUT2) caused a robust and rapid decrease in body temperature **(Figure 4**). Thermal imaging paired with photoactivation of terminals revealed that the decrease in body temperature was due to heat loss via rapid vasodilation and suppression of BAT thermogenesis (**Figure 4G-L**).

Previous studies have found that warmth activated PBN neurons were glutamatergic and projected to POA (Geerling et al., 2016). In the studies presented here, we found that hM4-Gi-DREADD mediated inhibition of VGLUT2+ (**Figure 5**) PBN neurons, which encompasses both the Enk and Dyn positive cells, was sufficient to block vasodilation in response to warm thermal challenge in awake animals. Activation of PBN→POA terminals leads to rapid vasodilation and hypothermia (**Figure 4**). Taken together, our results demonstrate the necessity and sufficiency of transmission from VGLUT2+ PBN neurons to the POA for physiological responses to thermal heat challenge.

In rodents, thermal heat stress evokes behavioral changes including grooming, suppression of physical activity, postural changes (postural extension), and thermal seeking (Roberts, 1988). Consistent with a neurobiological role in behavioral heat defense, photostimulation of Dyn^PBN→POA^ terminals markedly suppressed locomotor activity and evoked postural extension (**Figure 7**). The sprawled postural extension behavior is likely due to activation of cells in the POA by PBN terminals. Studies using lesion approaches in the POA have shown blocking of this behavior in response to warmth (Roberts and Martin, 1977), and selective activation of subpopulations of POA neurons can drive postural extension behavior (Yu et al., 2016). If subpopulations of POA cells independently drive individual behavioral components of thermal regulation such suppression of locomotion, postural extension, and alterations in metabolism will be an area for future investigation.

### Activation of PBN to POA projecting neurons drives avoidance but does not promote thermal cool seeking

The PBN and the POA have been found to play important roles in thermal seeking behaviors, but the neural circuitry involved remains poorly understood. Warmth activated neurons within the POA have previously been found to drive a temperature preference (Tan et al., 2016); however, the role of the POA in driving thermal seeking behaviors remains unclear. In contrast, prior studies using lesion approaches in the POA did not block thermal seeking behaviors (Almeida et al., 2006; Almeida et al., 2015; Matsuzaki et al., 2015). Studies examining the role of PBN in other aversive stimuli have found roles for the PBN in encoding valence and engaging motivational systems to drive avoidance without disruption of behaviors driven by sensory input. For example, functional silencing of LPBN Calcitonin gene-related peptide expressing neurons suppressed pain escape behavior; however, sensory reflex responses (paw withdrawal latency) remained intact (Han et al., 2015). In this example, disruption of PBN circuit activity blocks the expression of avoidance behaviors but not the transmission of sensory input.

The PBN may play a similar role, driving avoidance/escape behavior without altering sensation, in thermal defense. Muscimol mediated inhibition of PBN blocks temperature preference behavior, and a spinothalamic pathway independently conveys temperature information (Yahiro et al., 2017). We found that stimulation of PBN→POA terminals engages affective and motivational circuitry driving avoidance (**Figure 6**). Photoactivation of Dyn^PBN→POA^ terminals did not, however, induce a change in thermal preference for cooler temperatures **(Figure 7F-G).** In the context of previous studies, we interpret this to suggest that the coolness of the arena (20°C) as transduced sensory pathways remains aversive despite the decrease in body temperatures evoked by the same photostimulation. Taken together with the literature, the results presented here support the conclusion that PBN neurons are necessary, but activation of this pathway (PBN to POA) alone is not sufficient for expression of cold seeking behaviors. Thermal seeking may also require information from additional neural circuits, with the PBN encoding valence. An alternative is that additional targets of PBN neurons outside the POA may be required to engage thermal cool seeking behaviors, and those targets were not affected by our experimental photostimulation of POA terminal fields. Indicating that areas outside of the POA are required for thermal seeking, animals with POA lesion display amplification of motivated behaviors relating to thermal regulation due to impaired ability to defend core body temperature, and thus dependence on ambient temperature (Lipton, 1968; Satinoff et al., 1976). Future efforts will be necessary to better understand the select roles of POA neurons and PBN circuits in modulating thermal motivated behaviors.

### The endogenous opioid system is not required for acute effects of PBN neuron activation on body temperature

Opioid receptor modulation by agonists and antagonists has effects on body temperature regulation, acting at both central and peripheral sites and at mu, kappa and delta receptors (Baker and Meert, 2002). Specific effects of centrally administered mu and kappa antagonists on body temperature suggested a tonic balance between *kappa* and *mu* receptor activity in maintaining body temperature (Chen et al., 2005). Here we examined the potential roles of the endogenous opioid system in the acute hypothermic response evoked by activation of PBN→POA terminals in the POA by blocking opioid signaling with naloxone, naltrexone, or norBNI (**Supplemental Figure 3**). None of the selective opioid antagonists we used here significantly altered the response to acute stimulation of PBN terminals in the POA. One explanation for this lack of effect is that the PBN neuronal populations we examined are glutamatergic, and glutamate is known to be a key neurotransmitter for thermal regulation in the POA (Nakamura and Morrison, 2010). A role for the opioid system may be evoked by sustained changes in environmental temperature and may play a role in maintaining thermal set point in a more modulatory capacity. Additionally, our photo-activation paradigm might not be sufficient to produce endogenous opioid peptide release in these neurons. This is unlikely, however, given that our recent efforts in another region have shown that comparable photostimulation was sufficient to evoke both endogenous dynorphin and enkephalin release in vivo (Al-Hasani et al., 2018). Future studies with additional approaches and more sensitive peptide sensors may reveal further insights regarding the role of endogenous opioids in this circuitry.

### Future Directions and Conclusions

Previous studies have found that prior application of opioid receptor agonists affects the response of body temperature to opioid antagonists (Baker and Meert, 2002) and that environmental temperature, warm or cold, can dramatically alter the responses to centrally administered opioid peptides (Handler et al., 1994). Here we identified a potential source for multiple opioid peptides in the thermoregulatory neurocircuitry and delineated a role the neurons expressing Dyn and Enk in regulating body temperature. How these neuromodulators are involved in regulating body temperature and the target neurons will require further experimentation to delineate. How opioidergic circuits and signaling contribute to processes involving thermal regulation and dysregulations, such as during opiate withdrawal and alterations in calorie intake, merit further study. In sum, we report here that Dyn+, Enk+ and VGLUT2+ PBN neurons project to the POA, mediate physiological (vasodilation, suppression of thermogenesis) thermal defenses, drive behavioral thermal response behaviors (suppression of locomotion, postural changes), and drive aversion. The presented results will enable further studies to understand how homeostatic thermal regulation interacts with the motivational circuitry to drive behavior, provide targets for experiments testing the roles of neuromodulation of thermosensory pathways to regulate energy expenditure in balance with environmental factors, and help inform our understanding of how organisms balance competing interests, such as food intake, physical activity, and environmental conditions to in selecting behaviors.

## Acknowledgments

The authors thank Megan Votoupal for her technical assistance in mouse husbandry. This work was supported by a Foundation for Anesthesia Education and Research Grant, and Nation Institute for Mental Health grant K08MH119538 to AJN, by R01MH11235505 and R37DA03339607 to MRB, and by a Pilot Project Award from the Hope Center for Neurological Disorders at Washington University to AJN and MRB. The Mallinkrodt Foundation (MRB Professorship). The graphic abstract illustration was created by Percy Griffin with Astrid Rodriguez Velez in association with InPrint at Washington University School of Medicine.

## Author Contributions

Conceptualization, A.J.N., J.R.S, M.R.B.; Methodology, A.J.N., J.R.S; Investigation, A.J.N., J.R.S, A.L.C., I.B.N.; Writing –Original Draft, A.J.N., J.R.S; Writing –Review & Editing, A.J.N., M.R.B.; Visualization, A.J.N. J.R.S.; Funding Acquisition, A.J.N. and M.R.B.; Resources, A.J.N. and M.R.B..; Supervision, A.J.N. and M.R.B.

---

**Supplemental Figure 1.**
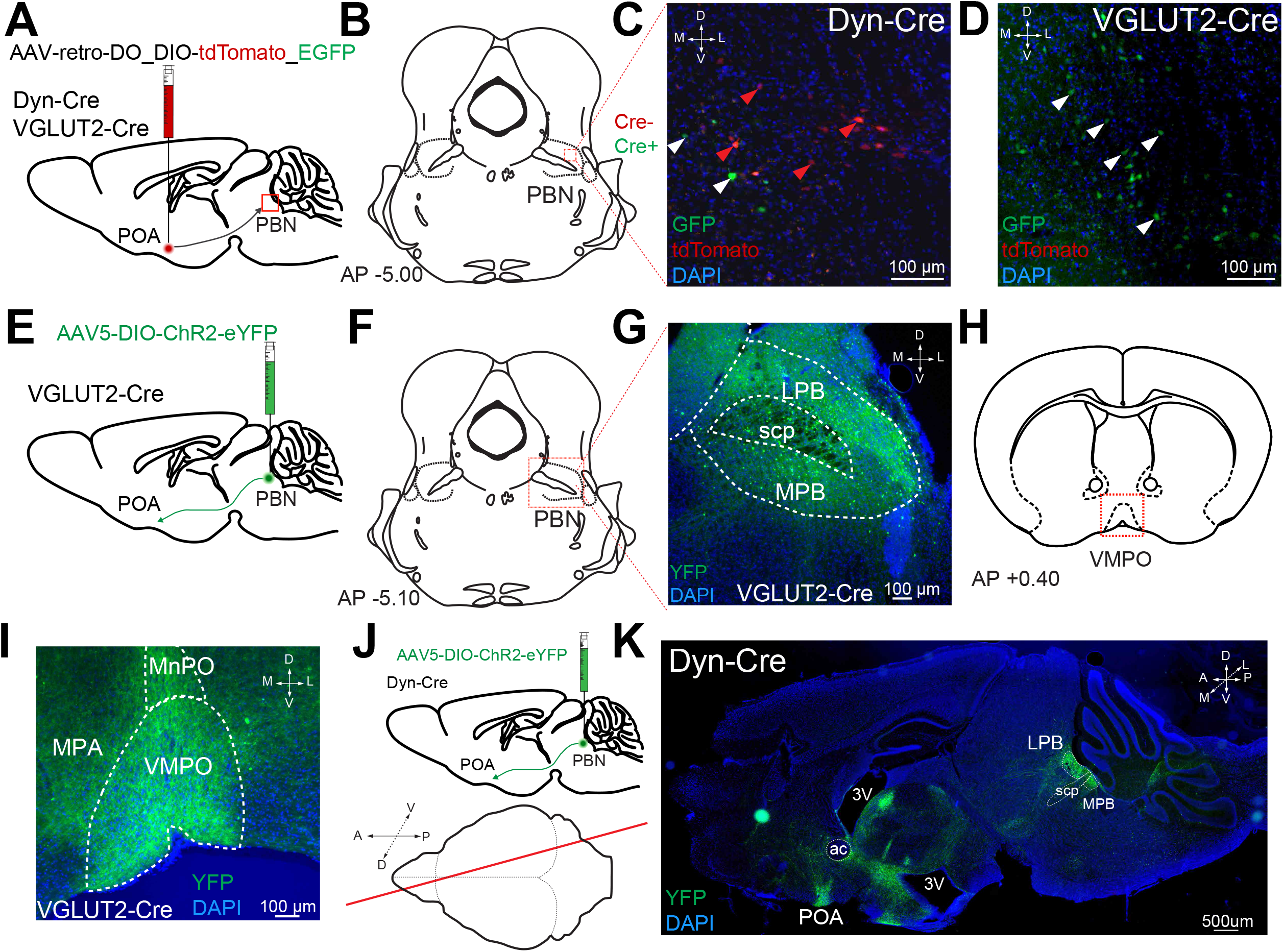
POA-projecting PBN neurons are VGLUT2+ and a subpopulation is Dyn+. Related to Figures 2 and 3. (**A**) Illustration of injections of a retrograde recombinase dependent red-to-green (tdTomato to EGFP) color changing virus (AAV-retro-DO_DIO-tdTomato_EGFP) into the POA of Dyn-Cre or VGLUT2-Cre mice. (**B**) Anatomical location of representative brain sections displayed in panels (C) (Dyn-Cre) and (D) (VGLUT2-Cre). Sections were analyzed for GFP and/or tdTomato expression in LPB. Cre-neurons transduced by the virus express tdTomato (red), Cre+ neurons express EGFP. (**C**) In brain sections taken from Dyn-Cre mice, neurons expressing tdTomato (Cre-, red) and EGFP (Cre+, green) are observed. White arrowheads mark EGFP expressing Cre+ cells; red arrowheads mark tdTomato expressing Cre-cells. (n=3 animals) (**D**) In brain sections taken from VGLUT2-Cre mice, only Cre+ cells expressing EGFP are observed (white arrowheads). (**E**) Illustration of viral injections into PBN of VGLUT2-Cre mice to label VGLUT2^PBN→POA^ projections. (**F**) Anatomical location of representative brain sections displayed in panel G. (**G**) Expression of eYFP in medial and lateral PBN of VGLUT2+ mice after injection of AAV5-DIO-ChR2eYFP into the PBN. (**H**) Anatomical location of representative brain sections displayed in panel (**I**) showing the POA projections of PBN VGLUT2+ neurons. (**J**) Illustration of injections into the PBN of Dyn-Cre mice to label anterograde projections and (lower) showing the plane of the sagittal section containing the PBN and the POA in (**K**) highlighting the dense projections of Dyn+ PBN neurons to the POA. (n=4 animals)

**Supplemental Figure 2.**
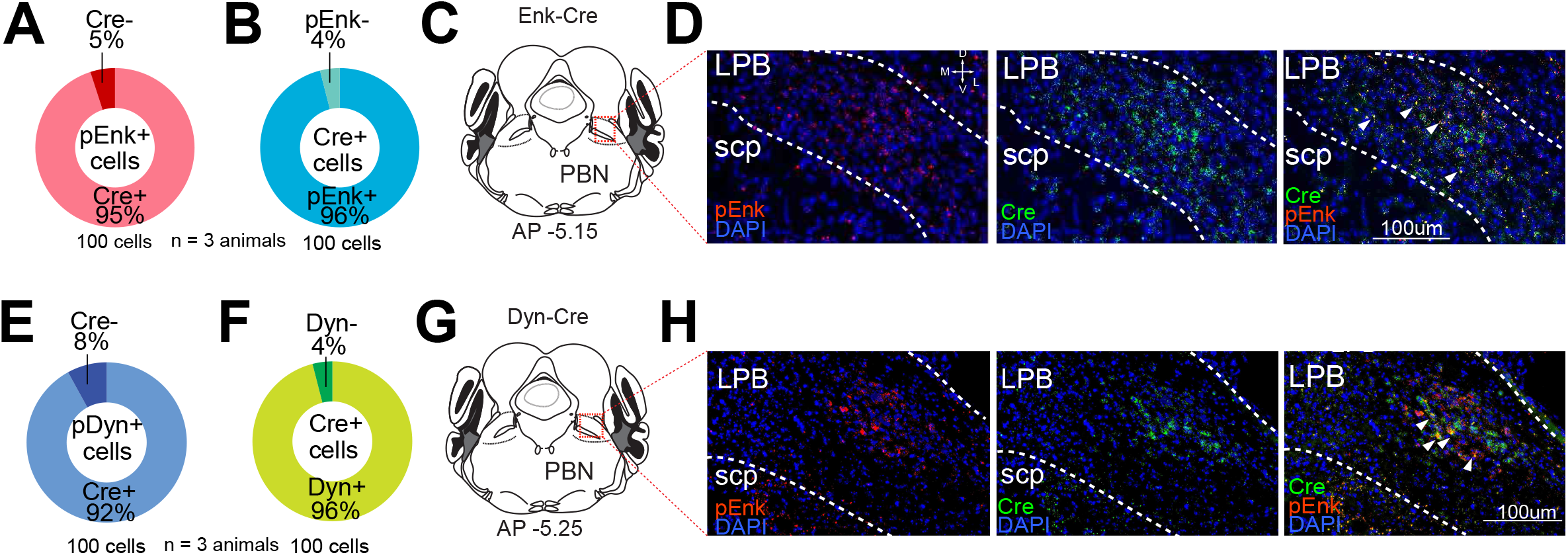
Validation of Enk-Cre and Dyn-Cre lines in the PBN. Related to Figures 1-3. FISH was performed on brain sections from Dyn-Cre or Enk-Cre mice using probes for prodynorphin or proenkephalin in combination with a probe for Cre transcripts. (**A-B**) The co-expression of Cre with proenkephalin was quantified by counting 100 cells from 3 animals positive for labeling of either (A) proenkephalin or (B) Cre and then counting the number of those cells that were also positive for the other probe. (**C**) Diagram of the anatomic localization of the PBN sections shown in (**D**) with labeling of proenkephalin (red), Cre (green) and DAPI (blue). Arrowheads highlight examples of cells labeled by probes for both transcripts. (**E-F**) The co-expression of Cre with prodynorphin was quantified by counting 100 cells from 3 animals positive for labeling of either (E) prodynorphin or (F) Cre and then counting the number of those cells that were also positive for the other probe. (**G**) Diagram of the anatomic localization of the PBN sections shown in (**H**) with labeling for prodynorphin (red), Cre (green) and DAPI (blue). Arrowheads highlight examples of cells labeled by probes for both transcripts.

**Supplemental Figure 3.**
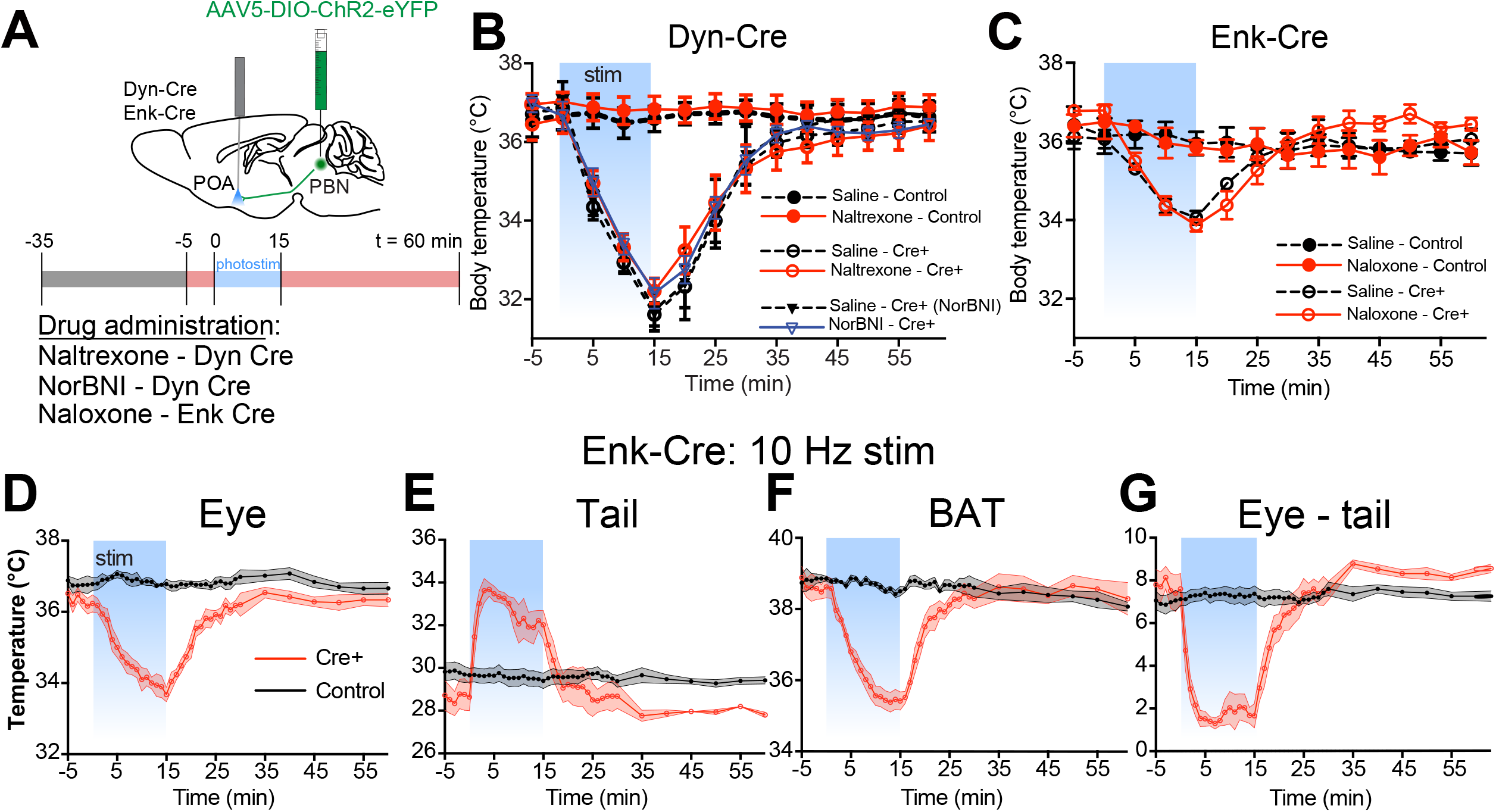
Dyn^PBN→POA^ or Enk^PBN→POA^ photostimulation-induced hypothermia is independent of opioid system. Related to Figure 4. (**A**) Illustrations of viral injections and fiber optic implantation in Dyn-Cre and Enk-Cre mice and timeline of drug administration/photostimulation/temperature recording trials. (**B**) Body temperature vs. time graphs for 10Hz photostimulation of Dyn^PBN**→**POA^ and control with IP saline, naltrexone (red) or NorBNI (blue) administration. Photostimulation was delivered from t=0 to t=15 min and led to equivalent levels of hypothermia in Cre+ mice regardless of pharmacologic pretreatment. Body temperature of controls was consistent throughout the trial. Data are presented as mean ± SEM. n=6-7 animals/group. (**C**) Body temperature vs. time graphs for 10Hz photostimulation of Enk^PBN**→**POA^ in Enk-Cre and control mice with prior IP naloxone or saline administration. Pretreatment did not alter the responses to photostimulation in Enk-Cre mice. Data are presented as mean ± SEM. n=5 animals in each group. (**D-G**) Results obtained using quantitative thermal imaging to measure tail, eye and BAT temperatures in Enk-Cre mice during 10 Hz photostimulation as done for Dyn-Cre mice in Figure 4. (**D**) Eye, (**E**) tail, (**F**) BAT, and (**G**) eye minus tail temperature vs. time graphs for 10 Hz photostimulation of Enk^PBN**→**POA^. Photostimulation was delivered from t=0 to t=15 min and led to drops in eye and BAT temperatures. Tail temperature rose and the gradient between eye and tail temperatures declined in Cre+ animals but not control mice. Data are presented as mean ± SEM. n=5 in each group.

**Supplemental Figure 4.**
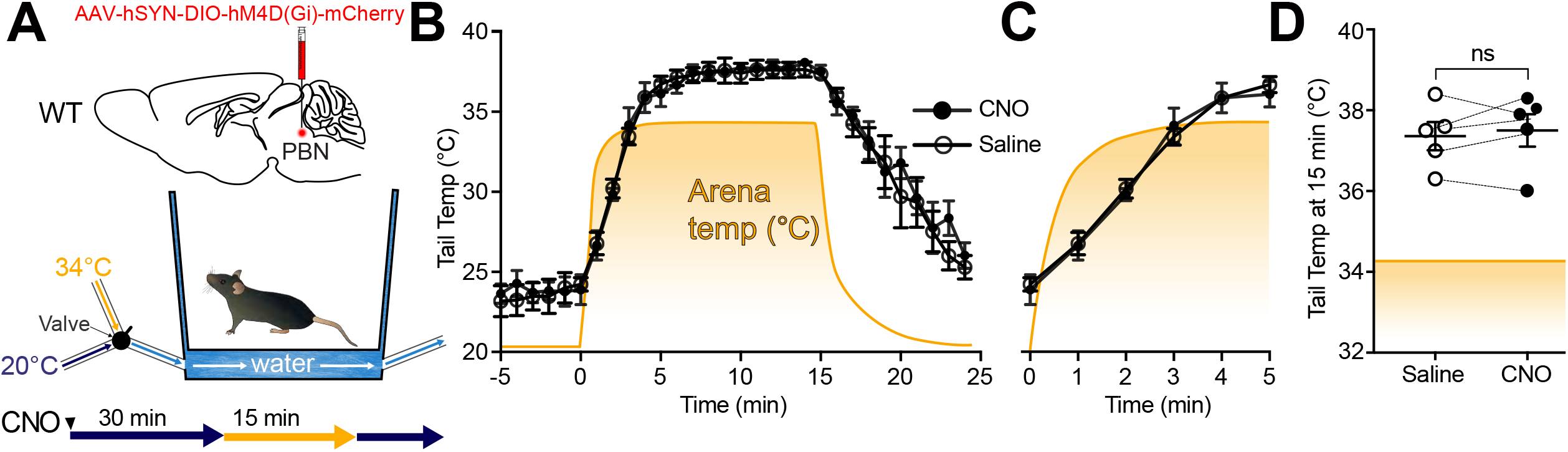
CNO does not alter responses to warmth challenge in wildtype mice. Related to Figure 5. (**A**) Illustrations depict viral injections in WT mice and arena that allowed for rapid changing of environmental temperature between two stable set points. (**B**) Tail temperature vs. time graph for 34°C heat challenge. Heat challenge was delivered from t=0 to t=15 min, and arena temperature throughout the trial is displayed by the orange line. In mice injected with CNO 2.5 mg/kg or in mice injected with saline, tail temperature rose above arena temperature after 4-5 minutes of heat challenge. Data are presented as mean ± SEM. n=5 animals, paired between CNO and saline conditions. (**C**) Tail temperature vs. time graph for 34°C heat challenge between t=0 and t=5 min. Note the overlap between average tail temperatures of the saline condition vs. the CNO condition across time. Data are presented as mean ± SEM. n=5 animals, paired between CNO and saline conditions. (**D**) Tail temperature at t=15 min of 34°C heat challenge. The average difference between tail temperatures in the saline condition and those in the CNO condition was not significantly different; 0.14±0.24°C. Paired t test, ns p = 0.59. Data are presented as mean ± SEM. n=5 animals, paired between CNO and saline conditions.

**Supplemental Figure 5.**
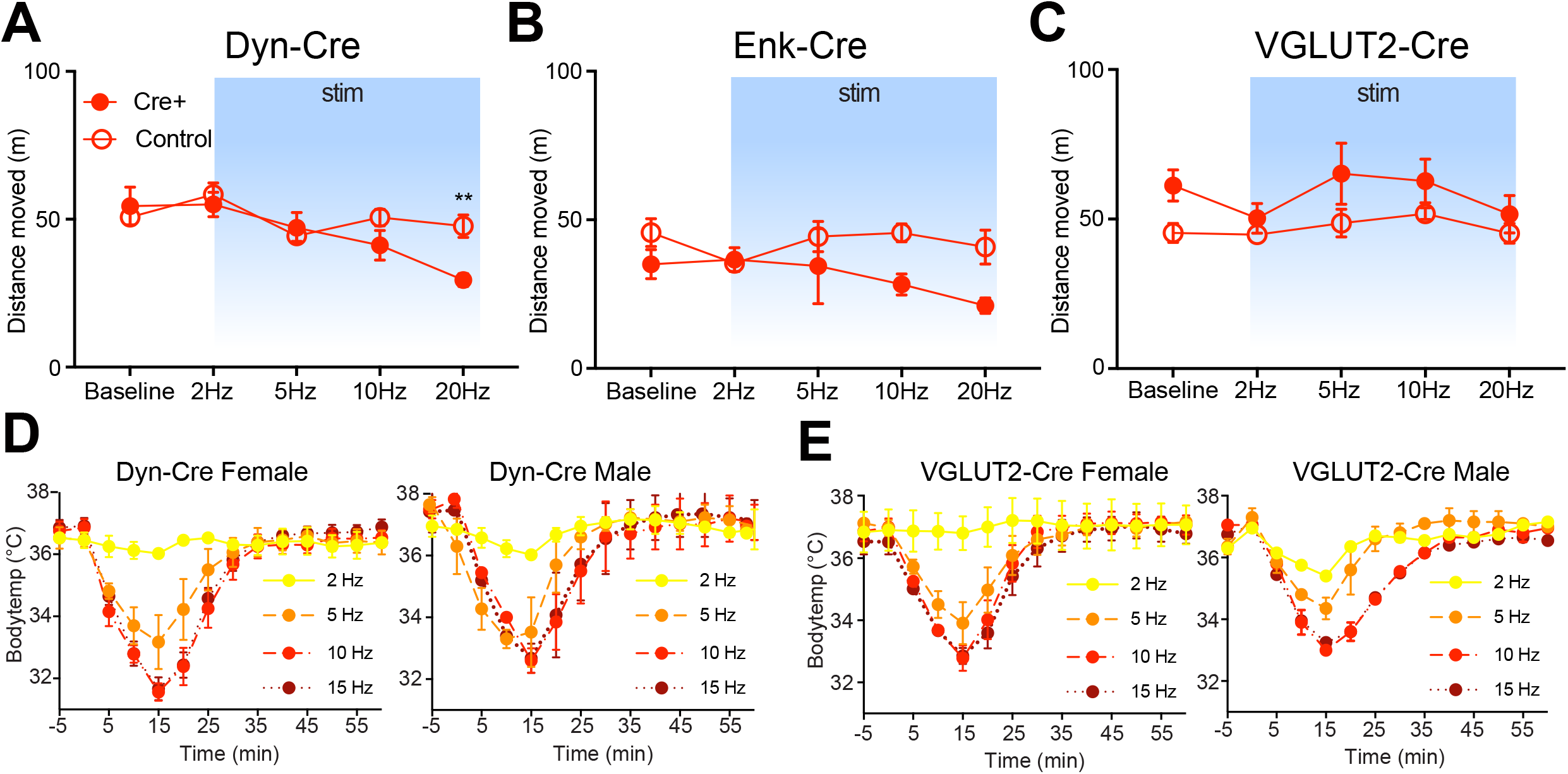
Total distance traveled for Dyn-Cre, Enk-Cre, and VGLUT2-Cre mice in Real-Time Place Aversion Assay and male vs. female photostimulation-induced body temperature change in Dyn-Cre and VGLUT2-Cre mice. Related to Figures 3 and 6. (**A**) Distance moved during RTPP trials displayed in **Figure 6C**(Dyn-Cre). Data are presented as mean ± SEM; n=6 Cre+, 8 control; two-way ANOVA, Bonferroni post hoc (20 Hz ChR2 vs. 20 Hz control **p < 0.01). (**B**) Distance moved during RTPP trials displayed in **Figure 6F**(Enk-Cre). Data are presented as mean ± SEM; n=6 Cre+, 7 control; two-Way ANOVA, Bonferroni post hoc (not significant at all frequencies ns p > 0.05). (**C**) Distance moved during RTPP trials displayed in **Figure 6I**(VGLUT2-Cre). Data are presented as mean ± SEM; n=8 Cre+, 7 control; two-Way ANOVA, Bonferroni post hoc (not significant at all frequencies ns p > 0.05). (**D-E**) The change in body temperature of Dyn-Cre female (n=4) and male (n=2) and VGLUT2-Cre female (n=6) and male (n=2) mice was similar at all stimulation frequencies (2, 5, 10, 15Hz) tested.

## STAR Methods

### Key Resources Table

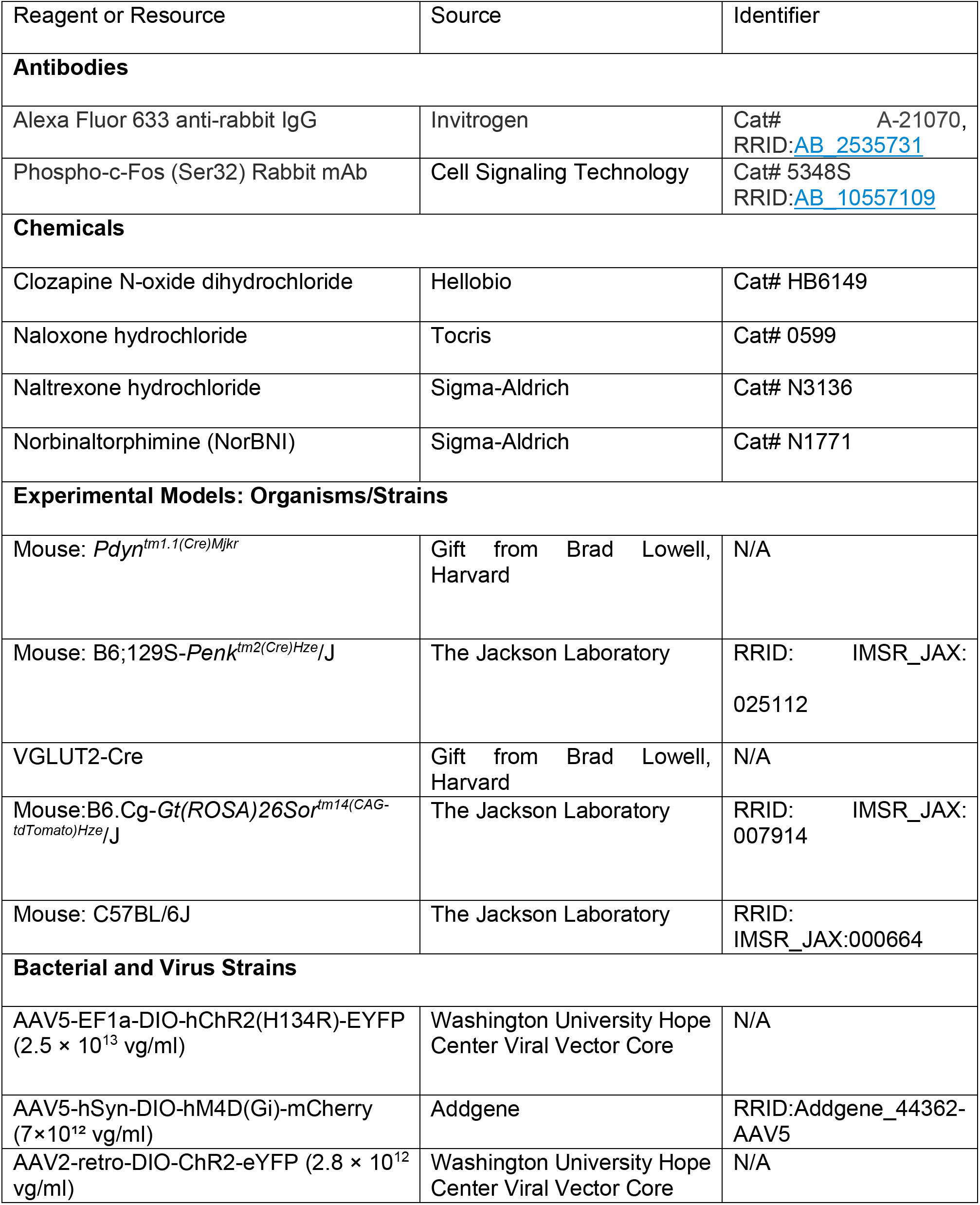

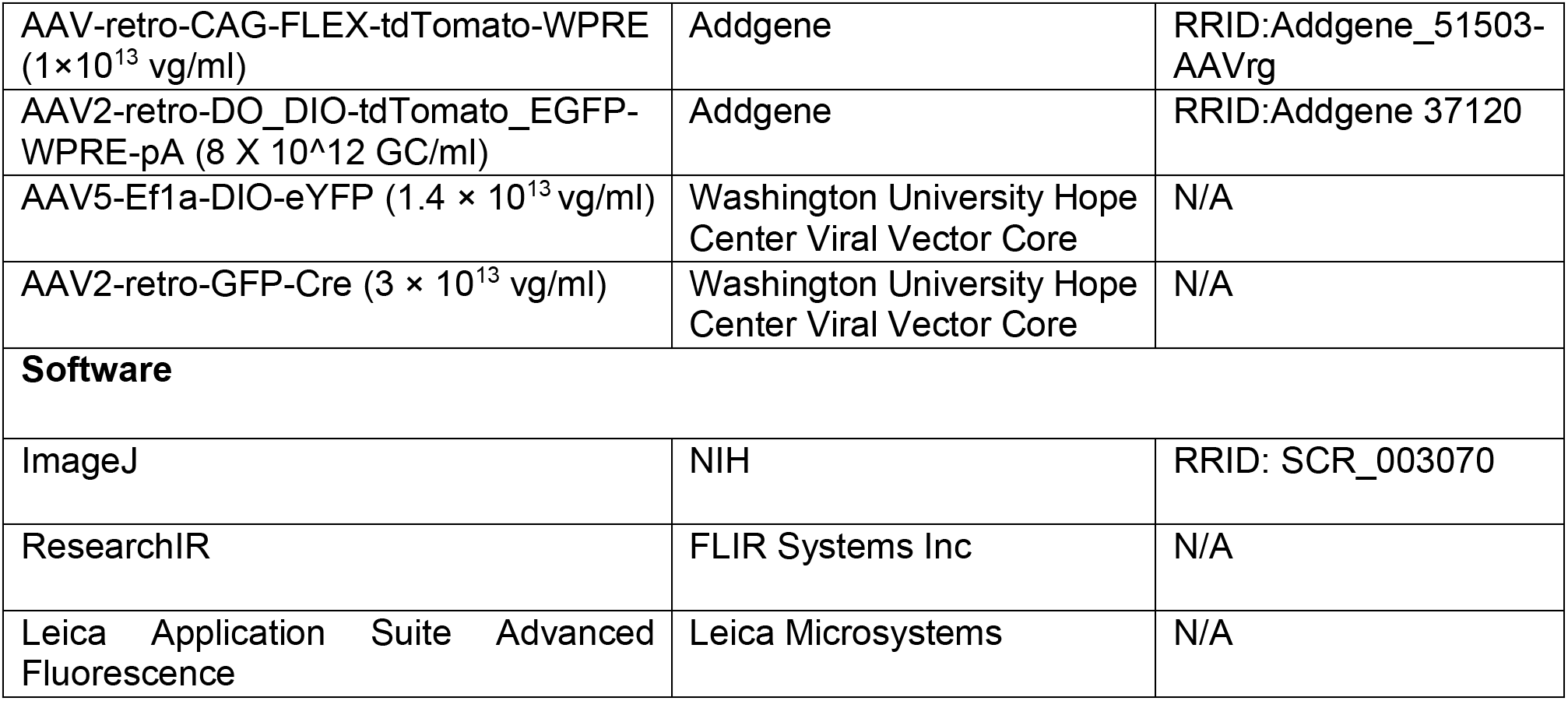

| Reagent or Resource | Source | Identifier |
| --- | --- | --- |
| **Antibodies** |  |  |
| Alexa Fluor 633 anti-rabbit IgG | Invitrogen | Cat# A-21070,<br>RRID:AB_2535731 |
| Phospho-c-Fos (Ser32) Rabbit mAb | Cell Signaling Technology | Cat# 5348S<br>RRID:AB_10557109 |
| **Chemicals** |  |  |
| Clozapine N-oxide dihydrochloride | Hellobio | Cat# HB6149 |
| Naloxone hydrochloride | Tocris | Cat# 0599 |
| Naltrexone hydrochloride | Sigma-Aldrich | Cat# N3136 |
| Norbinaltorphimine (NorBNI) | Sigma-Aldrich | Cat# N1771 |
| **Experimental Models: Organisms/Strains** |  |  |
| Mouse: *Pdyn*<sup>tm1.1(Cre)Mjkr</sup> | Gift from Brad Lowell, Harvard | N/A |
| Mouse: B6;129S-*Penk*<sup>tm2(Cre)Hze/J</sup> | The Jackson Laboratory | RRID: IMSR_JAX:025112 |
| VGLUT2-Cre | Gift from Brad Lowell, Harvard | N/A |
| Mouse: B6.Cg-*Gt(ROSA)26Sor*<sup>tm14(CAG-tdTomato)Hze/J</sup> | The Jackson Laboratory | RRID: IMSR_JAX:007914 |
| Mouse: C57BL/6J | The Jackson Laboratory | RRID: IMSR_JAX:000664 |
| **Bacterial and Virus Strains** |  |  |
| AAV5-EF1a-DIO-hChr2(H134R)-EYFP (2.5 × 10<sup>13</sup> vg/ml) | Washington University Hope Center Viral Vector Core | N/A |
| AAV5-hSyn-DIO-hM4D(Gi)-mCherry (7 × 10<sup>12</sup> vg/ml) | Addgene | RRID: Addgene_44362-AAV5 |
| AAV2-retro-DIO-ChR2-eYFP (2.8 × 10<sup>12</sup> vg/ml) | Washington University Hope Center Viral Vector Core | N/A |
| AAV-retro-CAG-FLEX-tdTomato-WPRE (1×10<sup>13</sup> vg/ml) | Addgene | RRID:Addgene_51503-AAVrg |
| AAV2-retro-DO_DIO-tdTomato_EGFP-WPRE-pA (8 X 10<sup>12</sup> GC/ml) | Addgene | RRID:Addgene 37120 |
| AAV5-Ef1a-DIO-eYFP (1.4 × 10<sup>13</sup> vg/ml) | Washington University Hope Center Viral Vector Core | N/A |
| AAV2-retro-GFP-Cre (3 × 10<sup>13</sup> vg/ml) | Washington University Hope Center Viral Vector Core | N/A |
| **Software** |  |  |
| ImageJ | NIH | RRID: SCR_003070 |
| ResearchIR | FLIR Systems Inc | N/A |
| Leica Application Suite Advanced Fluorescence | Leica Microsystems | N/A |

#### Contact for Reagent and Resource Sharing

Further information regarding reagents and resources may be directed to Aaron Norris.

#### Experimental Model and Subject Details

Adult (25-35 g, older than 8 weeks of age during experiments) male and female Dyn-Cre (Krashes et al., 2014), Enk-Cre (Harris et al., 2014), Ai14-tdTomato (Madisen et al., 2010), and VGLUT2-Cre (Vong et al., 2011) mice (species *Mus musculus*) were group housed (no more than 5 littermates per cage) and allowed food and water *ad libitum.* Mice were maintained on a 12 hr:12 hr light:dark cycle (lights on at 7:00 am). All procedures were approved by the Animal Care and Use Committee of Washington University and adhered to NIH guidelines. The mice were bred at Washington University in Saint Louis by crossing the Dyn-Cre, Enk-Cre, Ai14-tdTomato, and VGLUT2-Cre mice with C57BL/6 wild-type mice and backcrossed for seven generations. Additionally, where needed, Dyn-Cre and Enk-Cre mice were then crossed to Ai14-tdTomato mice on C57BL/6 background. Male and female mice were included and analyzed together.

#### Stereotaxic Surgery

Mice were anesthetized in an induction chamber (4% isoflurane), placed in a stereotaxic frame (Kopf Instruments), and anesthesia was maintained with 2% isoflurane. Mice were then injected bilaterally using a blunt needle Neuros Syringe (65457-01, Hamilton Com.) and syringe pump (World Precision Instruments) according to the injection schemes in the table below. The animal was kept in a warmed recovery chamber until recovery from anesthesia before being returned to its home cage.

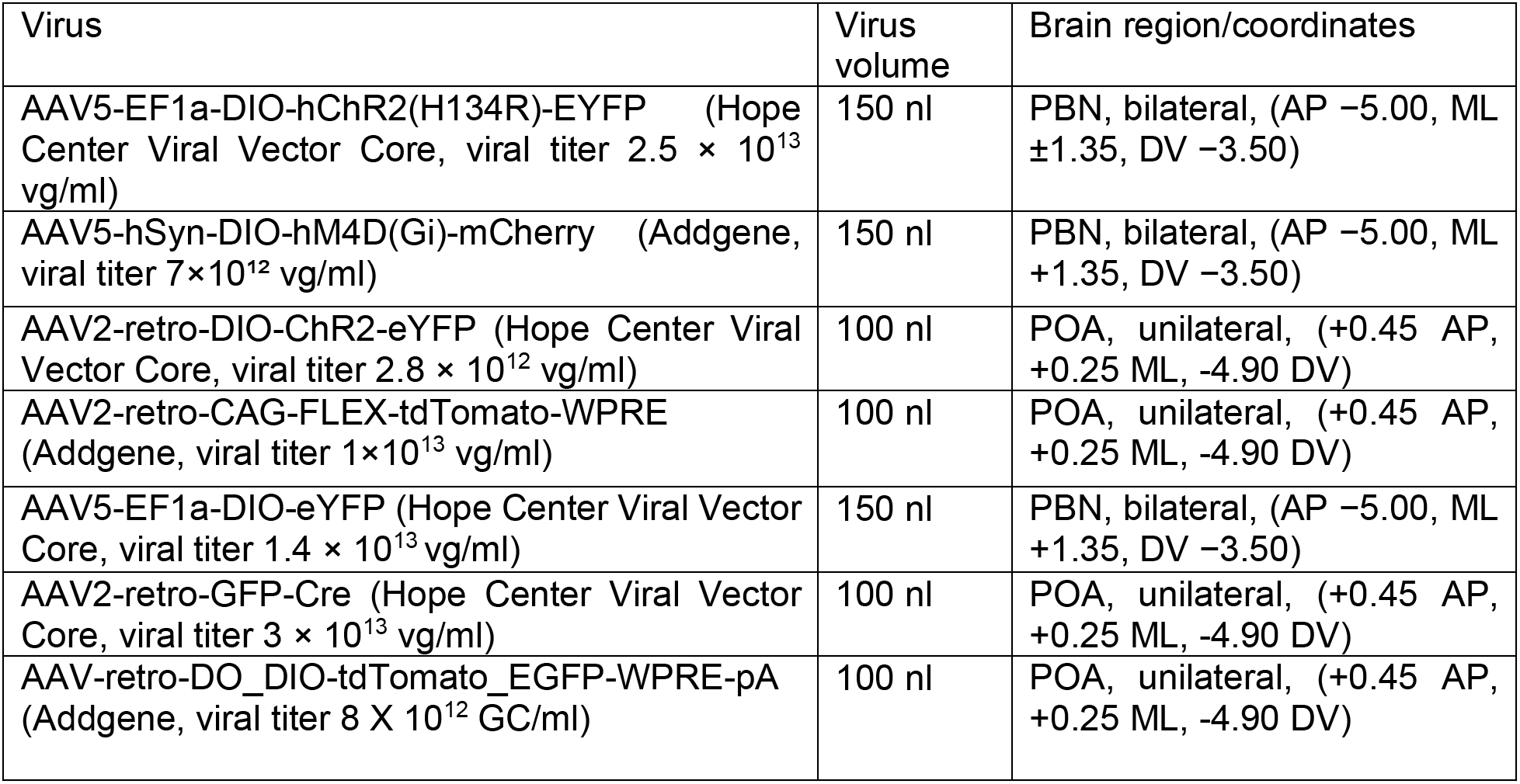

150 nl injections were injected at a rate of 30 nl/min, while 100 nl injections were injected at a rate of 20 nl/min. The injection needle was withdrawn 5 min after the end of the infusion. For anatomic experiments, mice that received unilateral or bilateral injections did not undergo further surgical procedures. For all behavioral experiments, mice underwent bilateral injections, implantations of a fiber optic for photostimulation over POA, and were implanted with a wireless IPTT-300 temperature transponder (Bio Medic Data Systems) subdermally directly rostral to right hindleg.

For photostimulation of PBN to POA projections, mice were injected with AAV5-EF1a-DIO-hChR2(H134R)-EYFP and were allowed 6 weeks for sufficient proteins to reach distal axons. Mice were then implanted with mono fiber optic cannulas (ChR2 mice: Thor Labs, 1.25 mm OD ceramic ferrule, 5 mm cannula with 200 μm OD, 0.22 NA) in the VMPO (+0.45 AP, +0.25 ML, and −4.60 DV for ChR2 mice). The fiber optic implants were affixed using Metabond (Parkell). Mice were allowed 7 days of recovery before the start of behavioral experiments. Viral injection coverage and optical fiber placements were confirmed in all animals using fluorescent microscopy in coronal sections (30 μm) to examine injection and implantation sites. Data from mice with incomplete viral coverage (i.e. unilateral expression of ChR2-eYFP in the PBN) or inaccurate optical fiber placement were excluded. Data from mice with bilateral PBN viral coverage and optical fiber placements near midline position over the POA were included in the study.

#### Anatomical Tracing

For anterograde viral tracing experiments, virus (AAV5-EF1a-DIO-hChR2(H134R)-EYFP or AAV5-EF1a-DIO-eYFP were used in our experiments) was injected at least six weeks prior to transcardial perfusions with 4% paraformaldehyde to allow for anterograde transport of the fluorophore. For retrograde viral tracing experiments, after the virus (AAV2-retro-DIO-ChR2-eYFP, AAV2-retro-CAG-FLEX-tdTomato-WPRE, AAV2-retro-EF1a-DO_DIO-TdTomato_EGFP-WPRE-pA, or AAV2-retro-GFP-Cre) was injected, there was a three week wait prior to perfusion to allow sufficient time for retrograde transport of the virus.

#### Warm Temperature Exposure

Ai14xDyn-Cre and Ai14xEnk-Cre mice in the warm condition were placed in a clean cage wrapped by a circulating water blanket which was set to 38°C. Mice in the room temperature condition were placed in a clean cage in a 22-23°C room. Water was supplied *ad libitum* in all cages. Cages in the warm condition were given enough time to reach the target temperature as confirmed by a thermometer before mice were placed inside of them. Temperature exposures lasted for 4 hours, after which mice were immediately anesthetized with pentobarbital and transcardially perfused with 4% paraformaldehyde in phosphate buffer, and brains were subsequently collected.

#### Immunohistochemistry

Immunohistochemistry was performed as previously described by (Al-Hasani et al., 2013, Kim et al., 2013, McCall et al., 2015). In brief, mice were intracardially perfused with 4% PFA and then brains were sectioned (30 microns) and placed in 1X PB until immunostaining. Free-floating sections were washed in 1X PBS for 3 ×10 min intervals. Sections were then placed in blocking buffer (0.5% Triton X-100 and 5% natural goat serum in 1X PBS) for 1 hr at room temperature. After blocking buffer, sections were placed in primary antibody (rabbit Phospho-c-Fos (Ser32) antibody (1:500), Cell Signaling Technology) overnight at room temperature. After 3 ×10 min 1X PBS washes, sections were incubated in secondary antibody (goat anti-rabbit Alexa Fluor 633 (1:1000), Invitrogen) for 2 hr at room temperature, followed by subsequent washes (3 × 10 min in 1X PBS then 3 × 10 min 1XPB washes). After immunostaining, sections were mounted on Super Frost Plus slides (Fisher) and covered with Vectashield Hard set mounting medium with DAPI (Vector Laboratories) and cover glass prior to being imaged on a Leica DM6 B microscope.

|  |  |  |  |  |
| --- | --- | --- | --- | --- |
| Alexa Fluor 633 anti-rabbit IgG | Goat | 1:1000 | Invitrogen | RRID:AB_2535731 |
| Phospho-c-Fos (Ser32) Rabbit mAb | Rabbit | 1:500 | Cell Signaling | RRID:AB_10557109 |

#### Imaging and Cell Quantification

Brain sections in figures are labeled relative to bregma using landmarks and neuroanatomical nomenclature as described in “The Mouse Brain in Stereotaxic Coordinates” (Franklin and Paxinos, 2013).

To quantify the number of cells expressing cFos, dynorphin, and/or enkephalin: cFos was labeled by Alexa Fluor 633, a fluorophore with emission in 610-800 nm (max 650 nm) range and preproenkephalin/prodynorphin were labeled by tdTomato with emission in the 540-700 nm (max 581 nm) range. All sections were imaged on a Leica DM6 B epifluorescent microscope using a Texas Red Filter Cube (Excitation: BP 560/40, Dichroic: LP 585, Emission: BP 630/75) for tdTomato visualization and a CY5 Filter Cube (Excitation: BP 620/60, Dichroic: LP 660, Emission: BP 700/75) for Alexa Fluor 633. Images were obtained for each 30 μm section that contained neurons in the PBN.

We defined the boundaries of LPB as follows. Sections between −5.0 and −5.4 rostral to bregma were imaged for LPB exclusively. The superior cerebellar peduncle marked the medial and ventral boundaries of LPB. The lateral boundary was marked by the ventral spinocerebellar tract, and the dorsal boundary was marked by the cuneiform nucleus.

All image groups were processed in parallel using ImageJ (RRID: SCR_003070, v1.50i) software. IHC was quantified as previously described (Al-Hasani et al., 2013; Kim et al., 2013b). Briefly, channels were separated, an exclusive threshold was set, and positive staining for each channel was counted in a blind-to-treatment fashion using ImageJ. The counts from each channel were then overlaid and percent of co-labeled cells were reported.

#### Fluorescent In Situ Hybridization (FISH)

Following rapid decapitation of mice, brains were flash frozen in −50°C 2-methylbutane and stored at −80°C for further processing. Coronal sections containing the PBN region, corresponding to the injection plane used in the behavioral experiments, were cut at 20μM at - 20°C and thaw-mounted onto Super Frost Plus slides (Fisher). Slides were stored at −80°C until further processing. FISH was performed according to the RNAScope® 2.0 Fluorescent Multiple Kit User Manual for Fresh Frozen Tissue (Advanced Cell Diagnostics, Inc.) as described by (Wang et al., 2012 - see below). Slides containing the specified coronal brain sections were fixed in 4% paraformaldehyde, dehydrated, and pretreated with protease IV solution for 30 mins. Sections were then incubated for target probes for mouse PDyn, (*Pdyn*, accession number NM_018863.3, probe region 33-700), PEnk (*Penk*, accession number NM_001002927.2, probe region 106-1332), VGLUT2 (slc17a6, accession number NM_080853.3, probe region 1986-2998), and/or Cre (accession number KC845567.1, probe region 1058-2032) for 2 hrs. All target probes consisted of 20 ZZ oligonucleotides and were obtained from Advanced Cell Diagnostics. Following probe hybridization, sections underwent a series of probe signal amplification steps (AMP1-4) including a final incubation of fluorescently labeled probes (Alexa 488, Atto 550, Atto 647), designed to target the specified channel (C1-C3 depending on assay) associated with the probes. Slides were counterstained with DAPI and coverslips were mounted with Vectashield Hard Set mounting medium (Vector Laboratories). Alternatively, mice transcardially perfused with cold PBS and PFA with fixed brain tissue collected and sectioned at 30 uM as described previously were processed for FISH as above.

Images were obtained on a Leica DM6 B upright microscope (Leica), and Application Suite Advanced Fluorescence (LAS AF) and ImageJ software were used for analyses. To analyze images for quantification of PDyn/PEnk/VGLUT2 coexpression, each image was opened in ImageJ software, channels were separated, and an exclusive fluorescence threshold was set. We counted total pixels of the fluorescent signal within the radius of DAPI nuclear staining, assuming that each pixel represents a single molecule of RNA as per manufacturer guidelines (RNAscope). A positive cell consisted of an area within the radius of a DAPI nuclear staining that measured at least 5 total positive pixels. Positive staining for each channel was counted in a blind-to-condition fashion using ImageJ or natively in LAX software (Leica).

#### Behavior

All behaviors were performed within a sound-attenuated room maintained at 23°C at least one week after the final surgery. For open field assays, lighting was stabilized at ∼250 lux for aversion behaviors (**Figures 6, 7, and S5A-C)** and ~200 lux for body temperature change recordings and heat challenges (**Figures 4, 5, S3, S4, and S5D-E**). Movements were video recorded and analyzed using Ethovision XT 10 (Noldus Information Technologies). For all optogenetic experiments, a 473 nm laser (Shanghai Lasers) was used and set to a power of ~15 mW from the tip of the patch cable ferrule sleeve. All patch cables used had a core diameter of 200 μm and a numerical aperture of 0.22 (Doric Lenses Inc.). At the end of each study, mice were perfused with 4% paraformaldehyde followed by anatomical analysis to confirm viral injection sites, optic fiber implant sites, and cell-type-specific expression.

#### Real-Time Place-Preference testing (RTPP)

We used 4 copies of a custom-made, unbiased, balanced two-compartment conditioning apparatus (52.5 × 25.5 × 25.5 cm) as described previously (Jennings et al., 2013; Stamatakis and Stuber, 2012). Mice were tethered to a patch cable and allowed to freely roam the entire apparatus for 30 min. Entry into one compartment triggered constant photostimulation at either 0 Hz (baseline trial), 2Hz, 5Hz, 10Hz, or 20 Hz (473 nm, 10 ms pulse width) while the mouse remained in the light paired chamber. Entry into the other chamber ended the photostimulation. The side paired with photostimulation was counterbalanced across mice. Ordering was counterbalanced with respect to stimulation frequency and placement in each of 4 of the copies of behavior apparatus. Bedding in all copies of the behavior apparatus was replaced between every trial, and the floors and walls of the apparatus were wiped down with 70% ethanol. Time spent in each chamber and total distance traveled for the entire 30-minute trial was measured using Ethovision 10 (Noldus Information Technologies).

#### Core Body Temperature, Vasodilation, and BAT Thermogenesis Recordings

We used transparent circular behavioral arenas (diameter = 14.5 inches, wall height = 21 cm) for experiments measuring core body temperature changes, vasodilation, and BAT thermogenesis suppression corresponding to optogenetic stimulation. Mice were tethered to a patch cable and allowed to habituate to the arena for 1 hour. Core body temperature measurements were made every 5 minutes beginning 5 minutes prior to turning the laser on. Laser frequencies of 2 Hz, 5 Hz, 10 Hz, and 15 Hz were used. Core body temperature measurements were made using a DAS-8007 Reader (Bio Medic Data Systems) which wirelessly read the temperature from a subdermally implanted IPTT-300 temperature transponder in each mouse (previously validated by (Langer and Fietz, 2014)).

Thermal imaging of mice was carried out using a FLIR E53 thermal imaging camera (FLIR Systems Inc) to record the 65-minute trial. Fur over the intrascapular region was shaven to facilitate temperature readings of the interscapular BAT(Crane et al., 2014). Thermal imaging videos were scored in a blind-to-genotype/condition fashion using ResearchIR software (FLIR Systems Inc). Eye, tail, and BAT temperatures were read every minute for the first 35 minutes of each trial and every 5 minutes for the final 30 minutes. Tail temperature readings were taken ~1 mm away from the base of the tail. BAT temperature readings were taken at the warmest point of the intrascapular region. Eye temperature readings were taken at the warmest point of the eye. To quantify the postural extension during these experiments, an investigator reviewed each video and quantified in two-minute bins the percent time the mice were in an extended posture (sprawled on the bedding) and time in huddled position with their tail tucked under their bodies.

#### Real-Time Place-Preference testing (RTPP)

We used 4 copies of a custom-made, unbiased, balanced two-compartment conditioning apparatus (52.5 × 25.5 × 25.5 cm) as described previously (Jennings et al., 2013; Stamatakis and Stuber, 2012). Mice were tethered to a patch cable and allowed to freely roam the entire apparatus for 30 min. Entry into one compartment triggered constant photostimulation at either 0 Hz (baseline trial), 2Hz, 5Hz, 10Hz, or 20 Hz (473 nm, 10 ms pulse width) while the mouse remained in the light paired chamber. Entry into the other chamber ended the photostimulation. The side paired with photostimulation was counterbalanced across mice. Ordering was counterbalanced with respect to stimulation frequency and placement in each of 4 of the copies of behavior apparatus. Bedding in all copies of the behavior apparatus was replaced between every trial, and the floors and walls of the apparatus were wiped down with 70% ethanol. Time spent in each chamber and total distance traveled for the entire 30-minute trial was measured using Ethovision 10 (Noldus Information Technologies).

#### Drug administration

Clozapine N-oxide dihydrochloride (Hellobio) was made in sterilized distilled water and mice received an intraperitoneal (i.p) injection of water (vehicle) or CNO (2.5 mg/kg) and were placed in the heat challenge apparatus or thermal plate preference apparatus for 30 minutes of habituation prior to beginning the assay. Naloxone hydrochloride (Tocris) and naltrexone hydrochloride (Sigma-Aldrich) were dissolved in 0.9% saline. Enk-Cre and Dyn-Cre mice received an i.p. injection of naloxone (5 mg/kg), naltrexone (3 mg/kg), or saline (vehicle) respectively and were placed back in their home cages for 30 minutes before being placed into behavioral arena. In experiments using NorBNI, NorBNI dissolved in DMSO (10mg/kg) was given IP approximately 24hrs prior to start of experiments and again 30mins immediately prior to start of the assay.

#### Heat Challenge

For chemogenetic inhibition experiments exposing mice to a heat challenge (**Figures 5 & S4**), we used a purpose-built, two-temperature water circulation apparatus to rapidly change the floor and wall temperatures of a square, transparent behavioral arena (15.25 × 15.25 × 19 cm). After drug or saline administration, mice were habituated to the arena at 20°C for 30 minutes. The water flow to the arena was changed to 34°C, and the temperature of floor and walls rose quickly, reaching steady state in the first 4 minutes (time course of ambient temperature change can be seen in **Figure 5B**). The water flow to the arena was switched back to 20°C after 15 minutes. Thermal imaging recording was obtained beginning 5 minutes prior to heat challenge and for 10 minutes post heat challenge for a total of 30 minutes. Thermal imaging videos were used to measure eye temperature, tail temperature, BAT temperature, and the temperature of the behavioral arena every minute throughout the 30-minute heat challenge trial. Thermal imaging videos were scored in a blind-to-genotype/condition fashion.

#### Thermal Preference

For experiments presenting mice with a choice between two floor plate temperatures (**Figure 7F-G**), we used a purpose-built apparatus consisting of 2 fused Cold / Hot Plate Analgesia Meters (Columbus Instruments International) with plastic walls surrounding and dividing the plates to create 2 behavioral arenas with 4-inch width, 19.5-inch length, and 9-inch height of walls. One side of the behavioral arena was set to 26°C and the other to 20°C. The side set to 20°C was counterbalanced across mice. Dyn-Cre mice were tethered to a patch cable and placed into a behavioral arena. Mice were allowed to roam the arena for 40 minutes before photostimulation. The laser frequency was set to 10 Hz and was left on for 20 minutes. Mice were kept in the behavioral arena for an additional 20 minutes post-stimulation. Time spent on each side for the entire 80-minute trial was quantified using Ethovision 10.

#### Open field test (OFT)

For experiments quantifying distance moved upon photostimulation of Dyn^PBN**→**POA^ (**Figure 7A-C**), we used a purpose-built 20in square behavior arena. Dyn-Cre mice were tethered to a patch cable and placed into the behavioral arena. The laser frequency was set to 10 Hz and was left on for 20 minutes. Distance moved for the 20-minute trial was quantified using Ethovision 10. Bedding in the arena was replaced between every trial, and the floors and walls of the arena were wiped down with 70% ethanol.

#### Statistical analyses

All data are expressed as mean ± SEM. Statistical significance was taken as *p < 0.05, **p < 0.01, and ***p < 0.001, ****p < 0.0001 as determined by student’s t-test, one-way ANOVA or a two-way repeated-measures ANOVA followed by a Bonferroni post hoc tests as appropriate. Statistical analyses were performed in GraphPad Prism 7.0. For each experiment, control groups and statistics are described in the main text. All “n” values represent the number of animals in a particular group for an experiment.

Experiments involving optogenetic stimulation of PBN inputs to POA using Dyn-Cre, Enk-Cre, and VGLUT2-Cre mice (**Figures 4, 6, 7, S3, and S5**) were replicated in 3 separate cohorts for each genotype. Chemogenetic inhibition experiments (**Figures 5 & S4**) were replicated in 2 separate cohorts of VGLUT2-cre mice and 2 separate cohorts of wild type mice. Warm temperature exposure with cFos immunohistochemical staining experiments (**Figure 1**) were performed in 4 separate iterations. Each iteration replicated the results of those prior to it, and data from each iteration was included in the overall statistical analysis of the experiment.

An investigator was blinded to allocation of groups in experiments whose data is shown in Figures 1 (warm-induced cFos+ cell quantification), 2 / S2 (in situ hybridization quantification), and 4 / 5 / 7 / S3 / S4 (thermal video scoring).

## References

Abbott, S.B.G., and Saper, C.B. (2017). Median preoptic glutamatergic neurons promote thermoregulatory heat loss and water consumption in mice. J Physiol 595, 6569–6583.

Abbott, S.B.G., and Saper, C.B. (2018). Role of the median preoptic nucleus in the autonomic response to heat-exposure. Temperature (Austin) 5, 4–6.

Al-Hasani, R., McCall, J.G., Shin, G., Gomez, A.M., Schmitz, G.P., Bernardi, J.M., Pyo, C.O., Park, S.I., Marcinkiewcz, C.M., Crowley, N.A., et al. (2015). Distinct Subpopulations of Nucleus Accumbens Dynorphin Neurons Drive Aversion and Reward. Neuron 87, 1063–1077.

Al-Hasani, R., Wong, J.T., Mabrouk, O.S., McCall, J.G., Schmitz, G.P., Porter-Stransky, K.A., Aragona, B.J., Kennedy, R.T., and Bruchas, M.R. (2018). In vivo detection of optically-evoked opioid peptide release. Elife 7.

Almeida, M.C., Steiner, A.A., Branco, L.G., and Romanovsky, A.A. (2006). Neural substrate of cold-seeking behavior in endotoxin shock. PLoS One 1, e1.

Almeida, M.C., Vizin, R.C., and Carrettiero, D.C. (2015). Current understanding on the neurophysiology of behavioral thermoregulation. Temperature (Austin) 2, 483–490.

Baker, A.K., and Meert, T.F. (2002). Functional effects of systemically administered agonists and antagonists of mu, delta, and kappa opioid receptor subtypes on body temperature in mice. J Pharmacol Exp Ther 302, 1253–1264.

Cabanac, M. (1975). Temperature regulation. Annu Rev Physiol 37, 415–439.

Chamberlin, N.L., Mansour, A., Watson, S.J., and Saper, C.B. (1999). Localization of mu-opioid receptors on amygdaloid projection neurons in the parabrachial nucleus of the rat. Brain Res 827, 198–204.

Chavkin, C., James, I.F., and Goldstein, A. (1982). Dynorphin is a specific endogenous ligand of the kappa opioid receptor. Science 215, 413–415.

Chen, X., McClatchy, D.B., Geller, E.B., Tallarida, R.J., and Adler, M.W. (2005). The dynamic relationship between mu and kappa opioid receptors in body temperature regulation. Life Sci 78, 329–333.

Chiang, M.C., Nguyen, E.K., Canto-Bustos, M., Papale, A.E., Oswald, A.M., and Ross, S.E. (2020). Divergent Neural Pathways Emanating from the Lateral Parabrachial Nucleus Mediate Distinct Components of the Pain Response. Neuron.

Cintron-Colon, R., Johnson, C.W., Montenegro-Burke, J.R., Guijas, C., Faulhaber, L., Sanchez-Alavez, M., Aguirre, C.A., Shankar, K., Singh, M., Galmozzi, A., et al. (2019). Activation of Kappa Opioid Receptor Regulates the Hypothermic Response to Calorie Restriction and Limits Body Weight Loss. Curr Biol 29, 4291–4299 e4294.

Clark, W.G. (1979). Influence of opioids on central thermoregulatory mechanisms. Pharmacol Biochem Behav 10, 609–613.

Crane, J.D., Mottillo, E.P., Farncombe, T.H., Morrison, K.M., and Steinberg, G.R. (2014). A standardized infrared imaging technique that specifically detects UCP1-mediated thermogenesis in vivo. Mol Metab 3, 490–494.

Czyzyk, T.A., Nogueiras, R., Lockwood, J.F., McKinzie, J.H., Coskun, T., Pintar, J.E., Hammond, C., Tschop, M.H., and Statnick, M.A. (2010). kappa-Opioid receptors control the metabolic response to a high-energy diet in mice. Faseb J 24, 1151–1159.

De Oliveira, L.B., De Luca, L.A., Jr., and Menani, J.V. (2008). Opioid activation in the lateral parabrachial nucleus induces hypertonic sodium intake. Neuroscience 155, 350–358.

Engstrom, L., Engblom, D., Ortegren, U., Mackerlova, L., Paues, J., and Blomqvist, A. (2001). Preproenkephalin mRNA expression in rat parabrachial neurons: relation to cells activated by systemic immune challenge. Neurosci Lett 316, 165–168.

Francois, A., Low, S.A., Sypek, E.I., Christensen, A.J., Sotoudeh, C., Beier, K.T., Ramakrishnan, C., Ritola, K.D., Sharif-Naeini, R., Deisseroth, K., et al. (2017). A Brainstem-Spinal Cord Inhibitory Circuit for Mechanical Pain Modulation by GABA and Enkephalins. Neuron 93, 822–839 e826.

Geerling, J.C., Kim, M., Mahoney, C.E., Abbott, S.B., Agostinelli, L.J., Garfield, A.S., Krashes, M.J., Lowell, B.B., and Scammell, T.E. (2016). Genetic identity of thermosensory relay neurons in the lateral parabrachial nucleus. Am J Physiol Regul Integr Comp Physiol 310, R41–54.

Gizowski, C., and Bourque, C.W. (2018). The neural basis of homeostatic and anticipatory thirst. Nat Rev Nephrol 14, 11–25.

Han, S., Soleiman, M.T., Soden, M.E., Zweifel, L.S., and Palmiter, R.D. (2015). Elucidating an Affective Pain Circuit that Creates a Threat Memory. Cell 162, 363–374.

Handler, C.M., Piliero, T.C., Geller, E.B., and Adler, M.W. (1994). Effect of ambient temperature on the ability of mu-, kappa- and delta-selective opioid agonists to modulate thermoregulatory mechanisms in the rat. J Pharmacol Exp Ther 268, 847–855.

Harris, J.A., Hirokawa, K.E., Sorensen, S.A., Gu, H., Mills, M., Ng, L.L., Bohn, P., Mortrud, M., Ouellette, B., Kidney, J., et al. (2014). Anatomical characterization of Cre driver mice for neural circuit mapping and manipulation. Front Neural Circuits 8, 76.

Henry, M.S., Gendron, L., Tremblay, M.E., and Drolet, G. (2017). Enkephalins: Endogenous Analgesics with an Emerging Role in Stress Resilience. Neural Plast 2017, 1546125.

Hermanson, O., and Blomqvist, A. (1997). Differential expression of the AP-1/CRE-binding proteins FOS and CREB in preproenkephalin mRNA-expressing neurons of the rat parabrachial nucleus after nociceptive stimulation. Brain Res Mol Brain Res 51, 188–196.

Hermanson, O., Telkov, M., Geijer, T., Hallbeck, M., and Blomqvist, A. (1998). Preprodynorphin mRNA-expressing neurones in the rat parabrachial nucleus: subnuclear localization, hypothalamic projections and colocalization with noxious-evoked fos-like immunoreactivity. Eur J Neurosci 10, 358–367.

Ikeda, T., Kurz, A., Sessler, D.I., Go, J., Kurz, M., Belani, K., Larson, M., Bjorksten, A.R., Dechert, M., and Christensen, R. (1997). The effect of opioids on thermoregulatory responses in humans and the special antishivering action of meperidine. Ann N Y Acad Sci 813, 792–798.

Jessen, C. (1985). Thermal afferents in the control of body temperature. Pharmacol Ther 28, 107–134.

Kim, D.Y., Heo, G., Kim, M., Kim, H., Jin, J.A., Kim, H.K., Jung, S., An, M., Ahn, B.H., Park, J.H., et al. (2020). A neural circuit mechanism for mechanosensory feedback control of ingestion. Nature 580, 376–380.

Kim, S.Y., Adhikari, A., Lee, S.Y., Marshel, J.H., Kim, C.K., Mallory, C.S., Lo, M., Pak, S., Mattis, J., Lim, B.K., et al. (2013). Diverging neural pathways assemble a behavioural state from separable features in anxiety. Nature 496, 219–223.

Krashes, M.J., Shah, B.P., Madara, J.C., Olson, D.P., Strochlic, D.E., Garfield, A.S., Vong, L., Pei, H., Watabe-Uchida, M., Uchida, N., et al. (2014). An excitatory paraventricular nucleus to AgRP neuron circuit that drives hunger. Nature 507, 238–242.

Langer, F., and Fietz, J. (2014). Ways to measure body temperature in the field. J Therm Biol 42, 46–51.

Lipton, J.M. (1968). Effects of Preoptic Lesions on Heat-Escape Responding and Colonic Temperature in Rat. Physiology & Behavior 3, 165−&.

Madisen, L., Zwingman, T.A., Sunkin, S.M., Oh, S.W., Zariwala, H.A., Gu, H., Ng, L.L., Palmiter, R.D., Hawrylycz, M.J., Jones, A.R., et al. (2010). A robust and high-throughput Cre reporting and characterization system for the whole mouse brain. Nat Neurosci 13, 133–140.

Mansour, A., Fox, C.A., Burke, S., Meng, F., Thompson, R.C., Akil, H., and Watson, S.J. (1994). Mu, delta, and kappa opioid receptor mRNA expression in the rat CNS: an in situ hybridization study. J Comp Neurol 350, 412–438.

Matsuzaki, K., Katakura, M., Sugimoto, N., Hara, T., Hashimoto, M., and Shido, O. (2015). beta-amyloid infusion into lateral ventricle alters behavioral thermoregulation and attenuates acquired heat tolerance in rats. Temperature (Austin) 2, 418–424.

Meng, F., Xie, G.X., Thompson, R.C., Mansour, A., Goldstein, A., Watson, S.J., and Akil, H. (1993). Cloning and pharmacological characterization of a rat kappa opioid receptor. Proc Natl Acad Sci U S A 90, 9954–9958.

Meyer, C.W., Ootsuka, Y., and Romanovsky, A.A. (2017). Body Temperature Measurements for Metabolic Phenotyping in Mice. Front Physiol 8, 520.

Miyaoka, Y., Shingai, T., Takahashi, Y., Nakamura, J.I., and Yamada, Y. (1998). Responses of neurons in the parabrachial region of the rat to electrical stimulation of the superior laryngeal nerve and chemical stimulation of the larynx. Brain Res Bull 45, 95–100.

Morrison, S.F. (2016). Central neural control of thermoregulation and brown adipose tissue. Auton Neurosci 196, 14–24.

Nakamura, K., and Morrison, S.F. (2007). Central efferent pathways mediating skin cooling-evoked sympathetic thermogenesis in brown adipose tissue. Am J Physiol Regul Integr Comp Physiol 292, R127–136.

Nakamura, K., and Morrison, S.F. (2010). A thermosensory pathway mediating heat-defense responses. Proc Natl Acad Sci U S A 107, 8848–8853.

Namburi, P., Al-Hasani, R., Calhoon, G.G., Bruchas, M.R., and Tye, K.M. (2016). Architectural Representation of Valence in the Limbic System. Neuropsychopharmacol 41, 1697–1715.

Palmiter, R.D. (2018). The Parabrachial Nucleus: CGRP Neurons Function as a General Alarm. Trends Neurosci 41, 280–293.

Qiu, M.H., Chen, M.C., Fuller, P.M., and Lu, J. (2016). Stimulation of the Pontine Parabrachial Nucleus Promotes Wakefulness via Extra-thalamic Forebrain Circuit Nodes. Curr Biol 26, 2301–2312.

Roberts, W.W. (1988). Differential thermosensor control of thermoregulatory grooming, locomotion, and relaxed postural extension. Ann N Y Acad Sci 525, 363–374.

Roberts, W.W., and Martin, J.R. (1977). Effects of lesions in central thermosensitive areas on thermoregulatory responses in rat. Physiol Behav 19, 503–511.

Satinoff, E., Valentino, D., and Teitelbaum, P. (1976). Thermoregulatory cold-defense deficits in rats with preoptic/anterior hypothalamic lesions. Brain Res Bull 1, 553–565.

Sheng, M., and Greenberg, M.E. (1990). The regulation and function of c-fos and other immediate early genes in the nervous system. Neuron 4, 477–485.

Siuda, E.R., Copits, B.A., Schmidt, M.J., Baird, M.A., Al-Hasani, R., Planer, W.J., Funderburk, S.C., McCall, J.G., Gereau, R.W.t., and Bruchas, M.R. (2015). Spatiotemporal control of opioid signaling and behavior. Neuron 86, 923–935.

Spencer, R.L., Hruby, V.J., and Burks, T.F. (1990). Alteration of thermoregulatory set point with opioid agonists. J Pharmacol Exp Ther 252, 696–705.

Stamatakis, A.M., and Stuber, G.D. (2012). Activation of lateral habenula inputs to the ventral midbrain promotes behavioral avoidance. Nat Neurosci 15, 1105–1107.

Tan, C.L., Cooke, E.K., Leib, D.E., Lin, Y.C., Daly, G.E., Zimmerman, C.A., and Knight, Z.A. (2016). Warm-Sensitive Neurons that Control Body Temperature. Cell 167, 47–59 e15.

Tan, C.L., and Knight, Z.A. (2018). Regulation of Body Temperature by the Nervous System. Neuron 98, 31–48.

Tan, K.R., Yvon, C., Turiault, M., Mirzabekov, J.J., Doehner, J., Labouebe, G., Deisseroth, K., Tye, K.M., and Luscher, C. (2012). GABA neurons of the VTA drive conditioned place aversion. Neuron 73, 1173–1183.

Vogel, B., Wagner, H., Gmoser, J., Worner, A., Loschberger, A., Peters, L., Frey, A., Hofmann, U., and Frantz, S. (2016). Touch-free measurement of body temperature using close-up thermography of the ocular surface. MethodsX 3, 407–416.

Vong, L., Ye, C.P., Yang, Z.F., Choi, B., Chua, S., and Lowell, B.B. (2011). Leptin Action on GABAergic Neurons Prevents Obesity and Reduces Inhibitory Tone to POMC Neurons. Neuron 71, 142–154.

Wolinsky, T.D., Carr, K.D., Hiller, J.M., and Simon, E.J. (1996). Chronic food restriction alters mu and kappa opioid receptor binding in the parabrachial nucleus of the rat: a quantitative autoradiographic study. Brain Res 706, 333–336.

Xin, L., Geller, E.B., and Adler, M.W. (1997). Body temperature and analgesic effects of selective mu and kappa opioid receptor agonists microdialyzed into rat brain. J Pharmacol Exp Ther 281, 499–507.

Yahiro, T., Kataoka, N., Nakamura, Y., and Nakamura, K. (2017). The lateral parabrachial nucleus, but not the thalamus, mediates thermosensory pathways for behavioural thermoregulation. Sci Rep 7, 5031.

Yu, S., Qualls-Creekmore, E., Rezai-Zadeh, K., Jiang, Y., Berthoud, H.R., Morrison, C.D., Derbenev, A.V., Zsombok, A., and Munzberg, H. (2016). Glutamatergic Preoptic Area Neurons That Express Leptin Receptors Drive Temperature-Dependent Body Weight Homeostasis. J Neurosci 36, 5034–5046.

